# Cryo-EM structures of α-synuclein fibrils with the H50Q hereditary mutation reveal new polymorphs

**DOI:** 10.1101/738450

**Authors:** David R. Boyer, Binsen Li, Chuanqi Sun, Weijia Fan, Michael R. Sawaya, Lin Jiang, David S. Eisenberg

**Affiliations:** Department of Chemistry and Biochemistry and Biological Chemistry, UCLA-DOE Institute, Molecular Biology Institute, and Howard Hughes Medical Institute, UCLA, Los Angeles CA 90095; Department of Neurology, David Geffen School of Medicine, Molecular Biology Institute, UCLA, Los Angeles, CA 90095 USA

## Abstract

Abstract

Deposits of amyloid fibrils of α-synuclein are the histological hallmarks of Parkinson’s disease, multiple system atrophy, and dementia with Lewy bodies. Although most cases of these diseases are sporadic, autosomal-dominant hereditary mutations have been linked to Parkinson’s disease and dementia with Lewy bodies. Seeing the changes to the structure of amyloid fibrils bearing these mutations may help to understand these diseases. To this end, we determined the cryo-EM structures of α-synuclein fibrils containing the H50Q hereditary mutation. We find that the H50Q mutation results in two new polymorphs of α-synuclein, which we term Narrow and Wide Fibrils. Both polymorphs recapitulate the conserved kernel formed by residues 50-77 observed in wild-type structures; however, the Narrow and Wide Fibrils reveal that H50Q disrupts a key interaction between H50-E57 on the opposing protofilament, abolishing the extensive protofilament interface formed by preNAC residues in the wild-type “rod” structure. Instead, the Narrow Fibril is formed from a single protofilament and the two protofilaments of the Wide protofilament are held together by only a pair of atoms – the Cɣ atoms from the two threonine 59 sidechains. Further, we find that H50Q forms an intramolecular hydrogen bond with K45 leading to the formation of a novel β-arch formed by residues 36-46 that features an extensive hydrogen-bond network between Y39, T44, and E46. The structures of the H50Q polymorphs help to rationalize the faster aggregation kinetics, higher seeding capacity in biosensor cells, and greater cytotoxicity we observe for H50Q compared to wild-type α-synuclein.

## Introduction

Several lines of evidence suggest that aggregation of α-synuclein (α-syn) into amyloid fibrils underlies the group of diseases termed synucleinopthies – Parkinson’s Disease (PD), Lewy Body Dementia (LBD), and Multiple Systems Atrophy (MSA): 1) α-syn fibrils are found in the hallmark lesions of PD and LDB – Lewy Bodies – as well as in the hallmark glial and neuronal lesions in MSA^1, 2^. 2) Hereditary mutations in α-syn have been linked to PD and LDB^3^. 3) Dominantly inherited duplications and triplications of the chromosomal region that contains wild-type *SNCA* – the gene that encodes α-syn – are sufficient to cause PD^4–6^. 4) Recombinantly assembled α-syn fibrils show cross-β structure and their injection into the brains of wild-type mice induced PD-like Lewy body and Lewy neurite formation, as well as cell-to-cell spreading, and motor deficits reminiscent of PD^7, 8^.

Advances in solid-state NMR and cryo-EM have greatly increased our knowledge of the structure of full-length amyloid proteins, allowing us to examine interactions beyond the local views provided by crystallographic methods^9–13^. Therefore, we previously used cryo-EM to determine the structures of wild-type full-length α-syn fibrils (Figure 1 b, left). These structures reveal two distinct polymorphs - termed the rod and twister^14^. Both fibrils are wound from two identical protofilaments related by an approximate 2_1_ fibril axis. The protofilaments that form the rod and twister polymorphs are distinct: the rod protofilaments contain ordered residues 38-97 whereas the twister protofilaments contain ordered residues 43-83. Both polymorphs share a similar structurally conserved β-arch formed by residues 50-77; however, the protofilament interfaces between the two polymorphs differ: in the rod polymorph residues 50-57 from the preNAC region form the interface of the two protofilaments whereas in the twister polymorph residues 66-78 from the NACore form the interface.

**Figure 1.**
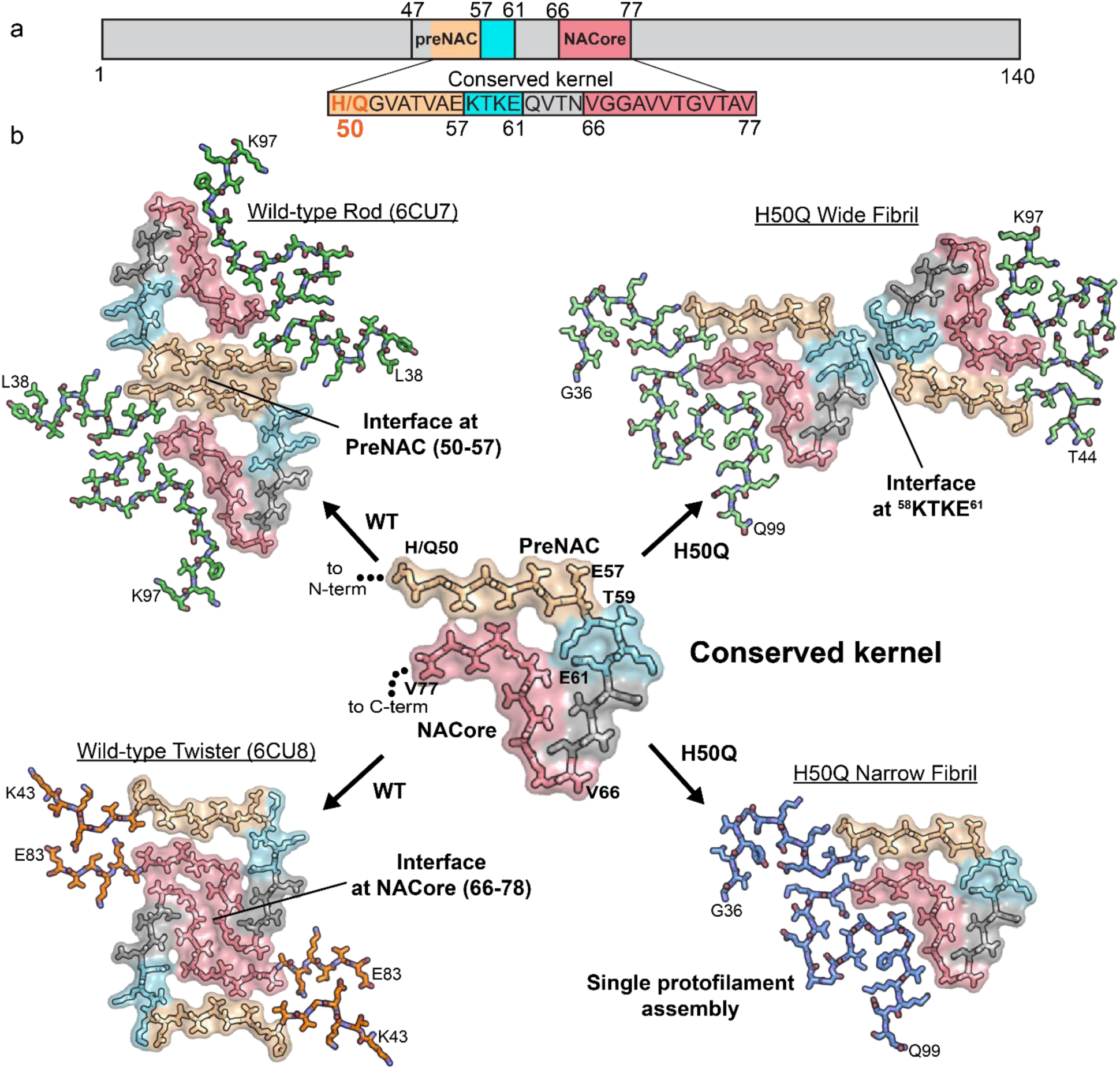
Comparison of wild-type and H50Q polymorphs. a) Primary structure schematic highlighting residues of the conserved kernel (50-77) that are used as protofilament interfaces in α-syn polymorphs. b) A conserved kernel is used to assemble wild-type (left) and H50Q (right) α-syn polymorphs in at least four different ways.

Hereditary mutations offer understanding of the link between protein structure and disease. H50Q is one such mutation that was discovered independently in two individuals with PD, with one patient having a known familial history of parkinsonism and dementia^15, 16^. The H50Q mutation enhances α-syn aggregation *in vitro* by reducing the solubility of monomer, decreasing the lag time of fibril formation, and increasing the amount of fibrils formed^17, 18^. Additionally, H50Q has been shown to be secreted at higher levels from SH-SY5Y cells and to be more cytotoxic to primary hippocampal neurons than wild-type α-syn^19^. Taken together these data suggest that patients harboring the H50Q mutation may develop fibrils more easily and that these fibrils may have different underlying structures than wild-type fibrils.

The structure of the wild-type rod polymorph suggests that the H50Q mutation may alter key contacts at the protofilament interface^14^. In this structure, two pairs of H50-E57 residues interact on opposing protofilaments stabilizing the protofilament interface through charge-charge interactions. The mutation to the uncharged, polar glutamine may therefore disrupt the rod polymorph protofilament interface, leading to different polymorphs of α-syn, potentially explaining the different observed properties of H50Q versus wild-type α-syn^17, 18^. In order to examine the exact effect of the H50Q mutation on the structure of α-syn fibrils, we sought to determine the atomic structures of H50Q α-syn fibrils using cryo-EM and to compare aggregation kinetics, stability, seeding capacity, and cytotoxicity to wild-type α-syn.

## Results

### Cryo-EM structure determination and architecture of H50Q α-syn fibrils

In order to pursue cryo-EM structure determination, we expressed and purified recombinant full-length α-syn containing the H50Q hereditary mutation and subsequently grew fibrils under the same conditions as those used for cryo-EM studies of wild-type α-syn (see *Methods*). After obtaining optimal cryo grid conditions and subsequent high-resolution cryo-EM data collection and processing, we determined the near-atomic structures of two polymorphs, which we term Narrow and Wide Fibrils, to resolutions of 3.3 and 3.6 Å, respectively (Figure 1 b, Figure 2 a-d, Supplementary Figure1 a-b, Supplementary Figure 2 a).

**Figure 2.**
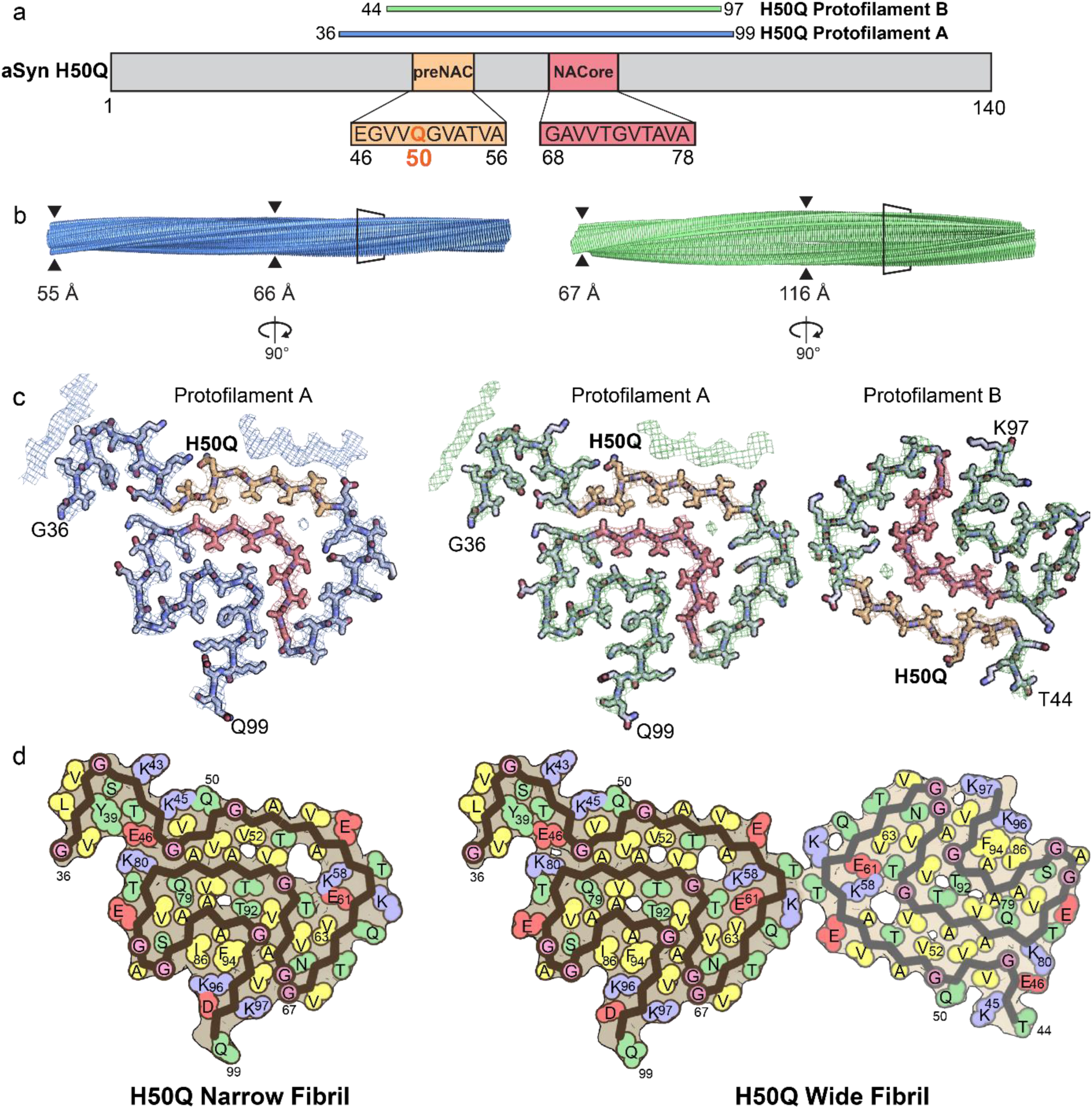
Cryo-EM Structures of H50Q Polymorphs a) Schematic of α-syn primary structure demonstrating location of the ordered core of H50Q Protofilaments A and B, preNAC and NACore, and H50Q hereditary mutation. b) View perpendicular to the fibril axes of Narrow and Wide Fibril cryo-EM reconstructions with the minimum and maximum widths of the fibrils labeled. c) View parallel to the fibril axes of sections of Narrow and Wide Fibrils revealing one layer of each fibril. Narrow Fibrils are composed of one protofilament designated Protofilament A. Wide Fibrils are composed of two protofilaments: Protofilament A and a less well ordered chain designated Protofilament B. Protofilament A is nearly identical in both fibril species while Protofilament B differs from Protofilament A and is only found in the Wide Fibril species. d) Schematic representation of fibril structures with amino acid side chains colored as follows: hydrophobic (yellow), negatively charged (red), positively charged (blue), polar, uncharged (green), and glycine (pink).

Both the Narrow and Wide Fibrils have pitches of ∼900 Å, as measured from the crossover distances observed in electron micrographs as well as 2D Classifications of box sizes containing helical segments that encompass an entire helical pitch (Supplementary Figure 3 a-b, see *Methods*). Narrow Fibrils have a width that varies from 55-66 Å; Wide Fibrils have a width that varies from 67-116 Å (Figure 2 b, Supplementary Figure 1 a). The Narrow Fibril is wound from a single protofilament, which we designate Protofilament A, while the Wide Fibril is wound from two slightly different protofilaments, both Protofilament A and a second protofilament which we term Protofilament B (Figure 2 c-d). Both Narrow and Wide Fibril reconstructions show identical densities flanking Protofilament A which we term Islands 1 and 2 (Supplementary Note 1, Supplementary Figure 4). Narrow Fibrils are roughly five times more abundant than Wide Fibrils. Protofilament A contains ordered residues from G36 to Q99 while Protofilament B contains ordered residues T44 to K97 (Fig. 2 a, c-d).

**Figure 3.**
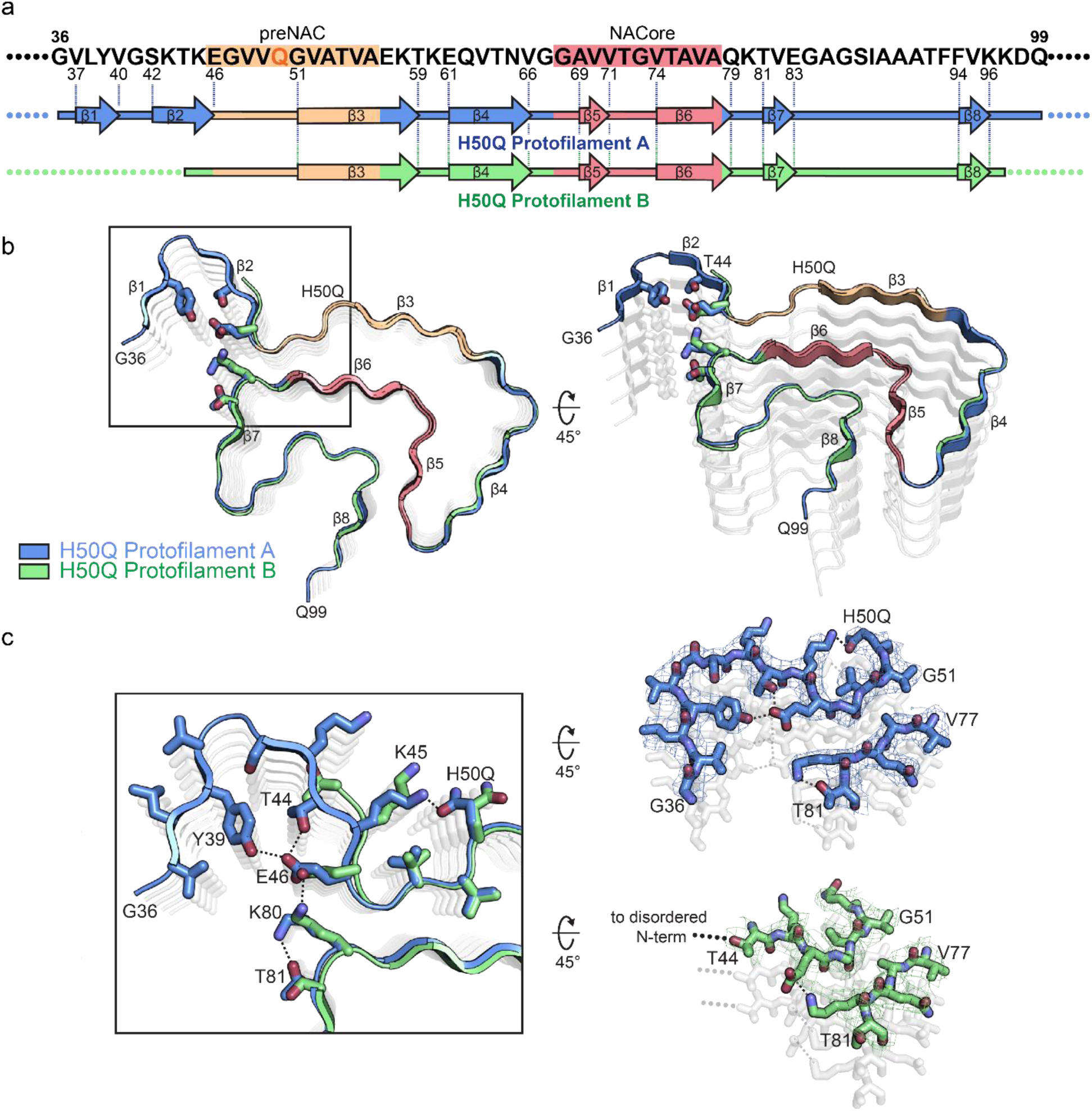
Comparison of Protofilaments A and B a) Schematic of primary and secondary structure of fibril core of H50Q Protofilaments A and B. Arrows indicate regions of Protofilaments A and B that adopt β-strand conformations. b) Structural alignment of H50Q Protofilaments A and B. c) Alignment of Protofilaments A and B reveals that Protofilament A has a β-arch formed by residues 36-46 whereas Protofilament B becomes disordered after residue 44. Additionally, in Protofilament A, H50Q hydrogen-bonds with K45 while in Protofilament B, H50Q and K45 adopt different conformations and do not hydrogen bond. In Protofilament A, E46 participates in the Y39-T44-E46 hydrogen bond triad and K80 hydrogen bonds with T81, while in Protofilament B E46 hydrogen bonds with K80 as in the wild-type rod polymorphs.

The protein chains in the ordered cores of both Protofilaments fold essentially within a two-dimensional layer (Figure 2 b-d), with stretches of straight β-strand regions interrupted by sharp turns (Figure 3 a-b). The fibrils are formed by the chains within the two-dimensional layers stacking upon one another along the fibril axis every 4.8 Å, forming β-sheets that extend for hundreds of nanometers (Figure 2 b).

### Differences between Protofilaments A and B

To define the differences of Protofilaments A and B observed in the two polymorphs, we aligned Protofilament A with Protofilament B (Figure 3 b). The alignment reveals that residues 47-97 adopt nearly identical conformations in both Protofilaments A and B; however, Protofilament A has an ordered β-arch formed by residues 36-46 whereas Protofilament B becomes disordered after T44 (Figure 3 b-c). The β-arch in Protofilament A features an extensive hydrogen bond network among Y39, T44, and E46 that is not observed in any α-syn structures determined to date (Figure 3 b-c). In addition, we note that in Protofilament A, Q50 is hydrogen bonded to K45 whereas in Protofilament B, K45 and Q50 are not hydrogen bonded and appear to be solvent-facing (Figure 3 b-c). This alternate arrangement of K45 and Q50 may explain why Protofilament A forms an ordered β-arch while Protofilament B does not. This suggests the conformation of K45 and Q50 can act as a switch: whereby as they become hydrogen-bonded, the N-terminal residues 36-44 become ordered, forming the Y39-T44-E46 hydrogen bond triad. We also note that Protofilament B maintains the E46-K80 hydrogen bond observed in wild-type structures while E46’s participation in a hydrogen bond with T44 in Protofilament A differs from wild-type structures. Instead, K80 now hydrogen bonds with T81 in Protofilament A (Figure 3 b-c). The differences observed here highlight the impact on fibril structure that hydrogen bonding arrangements can have and explain the atomic basis for the asymmetry of the two protofilaments in the Wide Fibril polymorph. Indeed, previous studies on wild-type α-syn fibril structures have revealed fibrils composed of asymmetric protofilaments whose asymmetry could be explained by the types of alternate hydrogen bond patterns we observe here^20^.

### Cavities in α-syn fibril structures

We note that all α-syn fibril structures determined to date display a similar cavity at the center of the β-arch surrounded by residues T54, A56, K58, G73, and V74 (Figure 1 b, Figure 2 c, Supplementary Figure 6 a, b). However, in contrast to our previously published wild-type structures, K58 now faces inward towards the cavity instead of outward towards the solvent, while T59 now flips away from the cavity (Supplementary Figure 6 a). This inversion is possible because lysine and threonine have both hydrophobic and hydrophilic character, allowing them to be favorably positioned either facing the solvent or the cavity. Indeed, energetic calculations demonstrate that K58 and T59 can have a positive stabilization energy in both cases (Supplementary Figure 5 b, c).

Presumably the β-arch cavity is filled with disordered solvent that is not defined by cryo-EM or ssNMR averaging methods. However, we note here that in our Narrow and Wide Fibril polymorphs, we visualize additional density in the cavity that is not accounted for by protein side chains (Figure 2 c, Supplementary Figure 6 b). We wonder whether this density arises from noise, or a back-projection artifact, but it is observed in the same location independently in three protofilaments (Protofilament A from the Narrow Fibril, and Protofilament A and B from the Wide Fibril) leading us to believe it may come from a solvent molecule (Supplementary Figure 6 b). Since our fibril growth conditions contain only water and tetrabutylphosphonium bromide, whose long aliphatic groups make it too large to fit into the tight cavity, we examined if the density could come from a water molecule and if any of the surrounding residues could serve as hydrogen bond partners. We observe that the γ-hydroxyl of T54, the ε-amino of K58, and the carbonyl oxygen of G73 are the only potential hydrogen bonding partners for the putative water molecule (Supplementary Figure 6 b). In addition, there are several methyl groups from V74 and A56 that are proximal to the putative solvent molecule (Supplementary Figure 6 b). The distance between the density and potential hydrogen partners ranges from 4.2 - 5.4 Å, longer than usual hydrogen bonds (Supplementary Figure 6 b). However, given that there are three hydrogen bond partners, perhaps the density visualized in the reconstruction is an average of positions occupied by water molecules. Nonetheless, given the resolution of our maps we cannot unambiguously identify the molecule(s) occupying the density in the center of the cavity.

### Wide Fibril Protofilament Interface

The Wide Fibril contains a novel protofilament interface formed by residues ^58^KTKE^61^ not previously seen in wild-type α-syn structures^14^. ^58^KTKE^61^ is located at a sharp turn in both Protofilaments A and B, and consequently the interface between protofilaments is remarkably small (Supplementary Figure 7 a). Consistent with the minimal size of the interface, the shape complementarity of 0.57 and buried surface area of 50 Å ^2^ of the Wide Fibril interface are low compared to the more extensive preNAC and NACore interfaces seen in our previous wild-type structures (Supplementary Figure 7 b)^14^. Indeed, the Cɣ atom of T59 from each protofilament is the only atom to interact across the protofilament interface (Supplementary Figure 7 b), making this interface the smallest fibril protofilament interface observed to date.

We wondered why the H50Q polymorphs do not utilize the extensive preNAC or NACore protofilament interfaces found in the wild-type rod and twister polymorphs, respectively. Our structures reveal that unlike the wild-type twister polymorph, the NACore is buried within the fibril core and is inaccessible as a protofilament interface (Figure 2 c). However, the preNAC region is more accessible as a potential protofilament interface in both the H50Q polymorphs (Figure 2 c). To examine why the preNAC region does not form a protofilament interface, we first compared the preNAC protofilament interface in the wild-type rod polymorph to the same region in the H50Q Narrow Fibril (Supplementary Figure 8 a-b). We noticed that in the wild-type rod polymorph H50 on one protofilament interacts with E57 on the opposite protofilament, possibly through a charge-charge interaction (Supplementary Figure 8 b). The mutation to a polar, uncharged glutamine leads to a loss of interaction with E57 and instead Q50 forms an intramolecular hydrogen bond with K45 in H50Q Protofilament A, thereby producing a single protofilament assembly (Supplementary Figure 8 a). Further, speculative models where additional protofilaments are docked at the preNAC region in either Protofilament A or B show large steric clashes occurring, further supporting the idea that the H50Q mutation disallows assembly of protofilaments at the preNAC (Supplementary Note 2, Supplementary Figure 8 c-d).

### Energetic and Biochemical Analysis of H50Q Fibrils

We next wondered if the structural differences we observe in the H50Q versus wild-type fibrils affect their stabilities. To examine this, we first calculated modified atomic solvation energies for the H50Q mutant structures and wild-type α-syn structures (see Supplementary Note 3 for differences in stabilization energies for some residues in Protofilaments A and B in the Wide Fibril). We find that both wild-type and H50Q fibrils are stabilized by energies comparable to a selection of known irreversible fibrils including tau Paired Helical Filament structures from Alzheimer’s Disease^21^, Serum Amyloid A fibrils from systemic amyloidosis^22^, and TDP-43 SegA-sym fibrils^23^ and significantly larger than the stabilization energies of the reversible FUS fibrils^24^ (Figure 4 a, Table 2). To confirm this, we performed stability assays to measure the resistance of fibrils to heat and SDS, and we found that both wild-type and H50Q show a similar resistance to denaturation, consistent with our energetic calculations (Figure 4 c). This is also consistent with the idea that both wild-type and H50Q α-syn are associated with pathogenic, irreversible fibrils formed in the synucleinopathies.

**Figure 4.**
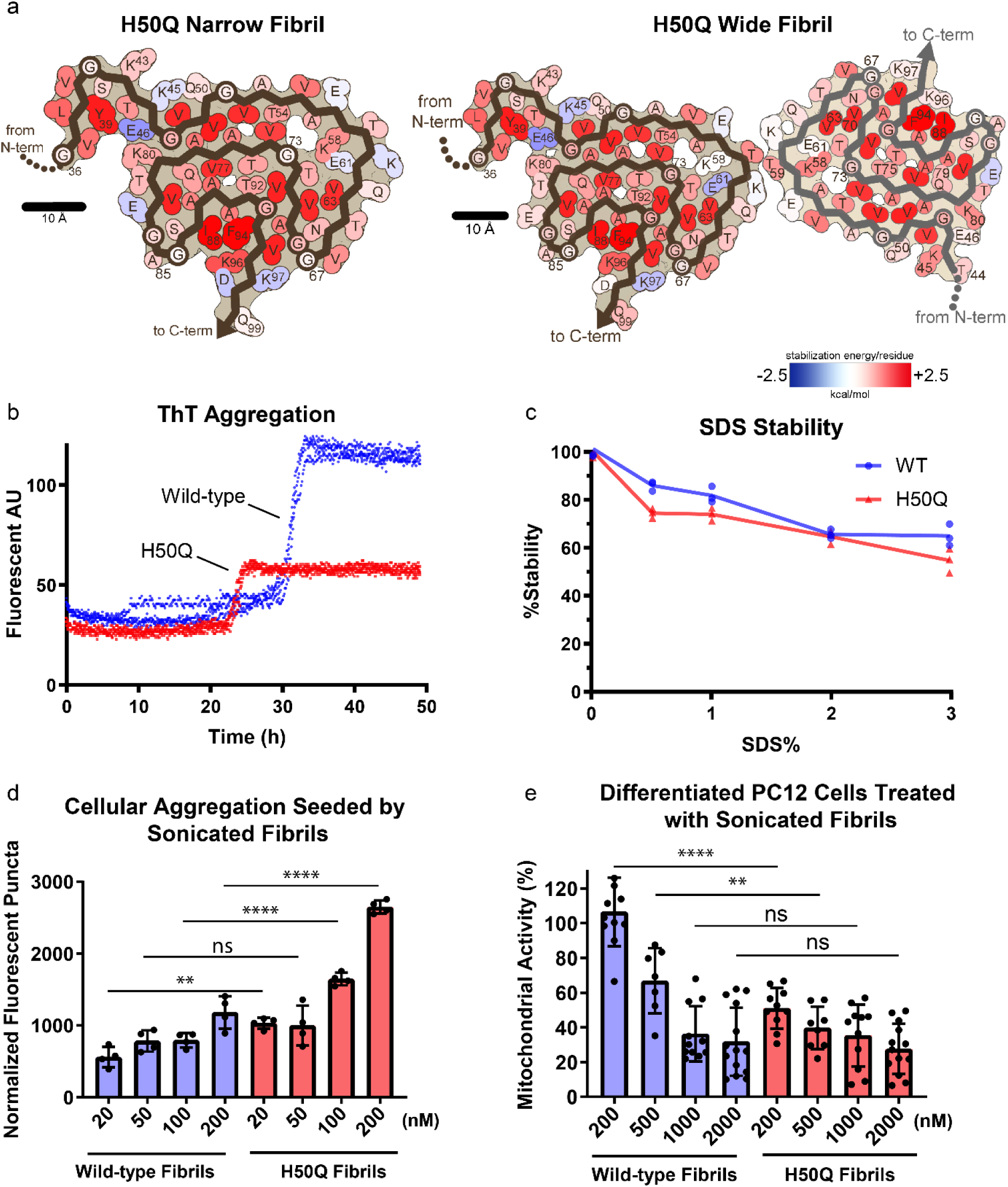
Solvation energy maps and biochemical characterization of H50Q and wild-type α- syn. a) Solvation energy maps of H50Q Narrow and Wide Fibrils. The stabilizing residues are red; the de-stabilizing residues are blue. b) ThT assay measuring kinetics of H50Q and wild-type α-syn aggregation. Data from three independent replicates for each sample are shown for each time point. Wild-type aggregation plateaus at 30 h whereas H50Q aggregates faster and plateaus at 23 h. c) Stability assay of H50Q and wild-type α-syn. Fibrils were brought to the indicated concentration of SDS and heated at 70° C for 15 minutes before ThT signal was measured. Both H50Q and wild-type fibrils are irreversible. d) Cell seeding assay of H50Q and wild-type α-syn performed in HEK293T α-syn A53T-YFP biosensor cells. Sonicated fibrils were transfected into biosensor cells using Lipofectamine. After 48 hrs., the number of fluorescent puncta indicating aggregated endogenous α-syn A53T-YFP were counted (see *Methods*). H50Q fibrils have a higher seeding capacity than wild-type fibrils. Error bars represent standard deviation of four independent measurements. e) MTT toxicity assay of H50Q and wild-type α-syn performed in differentiated PC12 cells. H50Q and wild-type α-syn fibrils were introduced to culture medium and after incubation, cell mitochondrial activity was measured via MTT assay. H50Q requires less fibrils than wild-type to significantly disrupt mitochondrial activity. Error bars represent standard deviation of a minimum of 7 and a maximum of 14 independent measurements. **** = p-value ≤ 0.0001. ** = p-value ≤ 0.01. ns = p-value > 0.05. P-values were calculated using an unpaired, two-tailed t-test with a 95% CI.

**Table 1.** Cryo-EM data collection, refinement, and validation statistics.

| Name | H50Q Narrow Fibril | H50Q Wide Fibril |
| --- | --- | --- |
| PDB ID | 6PEO | 6PES |
| EMDB ID | EMD-20328 | EMD-20331 |
| **Data collection** |  |  |
| Magnification | ×130,000 | ×130,000 |
| Defocus range (um) | 1.00-3.04 | 0.80-2.91 |
| Voltage (kV) | 300 | 300 |
| Camera | K2 Summit<br>(Quantum LS) | K2 Summit<br>(Quantum LS) |
| Frame exposure time (s) | 0.2 | 0.2 |
| # movie frames | 30 | 30 |
| Total electron dose (e<sup>-</sup>/ Å<sup>2</sup>) | 26 | 26 |
| Pixel size (Å) | 1.065 | 1.065 |
| **Reconstruction** |  |  |
| Box size (pixel) | 288 | 224 |
| Inter-box distance (Å) | 28.8 | 22.4 |
| # micrograph collected | 3,577 | 3,577 |
| # segments extracted | 1,183,284 | 137,395 |
| # segments after Class2D | 510,477 |  |
| # segments after Class3D | 30,133 | 28,016 |
| Resolution (Å) | 3.3 | 3.6 |
| Map sharpening B-factor (Å<sup>2</sup>) | -148 | -207 |
| Helical rise (Å) | 4.81 | 4.82 |
| Helical twist (°) | -0.97 | -0.83 |
| Point Group | C<sub>1</sub> | C<sub>1</sub> |
| **Atomic model** |  |  |
| # non-hydrogen atoms | 2205 | 4040 |
| # protein residues | 320 | 590 |
| R.m.s.d. bonds (Å) | 0.003 | 0.009 |
| R.m.s.d. angles (°) | 0.543 | 0.704 |
| Molprobit clashscore, all atoms | 10.62 | 12.28 |
| Molprobit score | 2.09 | 2.18 |
| Poor rotamers (%) | 0 | 0 |
| Ramachandran outliers (%) | 0 | 0 |
| Ramachandran allowed (%) | 9.7 | 12.3 |
| Ramachandran favoured (%) | 90.3 | 87.7 |
| Cβ deviations > 0.25 Å (%) | 0 | 0 |
| Bad bonds (%) | 0 | 0 |
| Bad angles (%) | 0 | 0 |

**Table 2.** Comparative solvation energy calculations.

| Fibril Structure | Atomic solvation standard free energy of stabilization<br>(kcal/mol) |  |  |
| --- | --- | --- | --- |
|  | Per Layer | Per Chain | Per Residue |
| H50Q Narrow Fibril | 58 | 58 (Protofilament A) | 0.90 (Protofilament A) |
| H50Q Wide Fibril | 114 | 58 (Protofilament A)<br>54 (Protofilament B) | 0.90 (Protofilament A)<br>1.0 (Protofilament B) |
| WT Rod CryoEM (6CU7) | 112 | 56 | 0.93 |
| WT Twister CryoEM (6CU8) | 66 | 33 | 0.80 |
| WT ssNMR (2N0A) | 58 | 58 | 0.91 |
| WT Rod (6H6B) | 101 | 51 | 0.86 |
| WT Rod (6A6B) | 122 | 61 | 0.96 |
| Tau PHF (530L) | 128 | 64 | 0.88 |
| Tau Pick's (6GX5) | 98 | 98 | 1.0 |
| TDP-43 SegA-sym (6N37) | 73 | 36.5 | 1.0 |
| Serum Amyloid A (6MST) | 116 | 58 | 1.07 |
| FUS (5W3N) | 41 | 41 | 0.66 |

We next characterized the kinetics of H50Q and wild-type fibril growth and found that H50Q aggregation has a shorter lag phase and therefore forms fibrils more rapidly than wild-type fibrils, consistent with other studies (Figure 4 c)^18, 19, 25^. We note that although H50Q fibrils have a lower max ThT signal, this is likely explained by different fibril polymorphs having differential ThT binding and not due to overall less fibril formation. We also find that H50Q fibrils have significantly higher seeding capacity in HEK293T α-syn-A53T-YFP biosensor cells^26^ and significantly higher cytotoxicity to differentiated neuron-like rat pheochromocytoma (PC12) as measured by a reduction of mitochondrial activity and cell membrane integrity (Figure 4 d-e, Supplementary Figure 9 a). These results are similar to other studies demonstrating the enhanced pathogenic properties of H50Q vs. WT α-syn^17–19, 25^, and overall our findings support the idea that the differences in the structures of the H50Q compared to wild-type fibrils result in higher pathogenicity.

## Discussion

The structures of α-syn fibrils determined here – the first atomic structures of α-syn fibrils with a hereditary mutation – demonstrate that the H50Q mutation results in two new polymorphs, which we term Narrow and Wide Fibrils. Structural alignments of wild-type and H50Q polymorphs reveal that residues 50-77 in all structures adopt a largely similar β-arch-like fold, which we previously termed the conserved kernel (Supplementary Figure 10)^14^. However, different sequence segments of the conserved kernel assemble protofilaments into distinct fibril polymorphs (Figure 1 a-b). The wild-type rod polymorph utilizes residues from the preNAC region^14, 27, 28^, the wild-type twister polymorph utilizes residues from the NACore^14^, and the H50Q Wide Fibril utilizes residues ^58^KTKE^61^ to assemble protofilaments while the H50Q Narrow Fibril forms a single protofilament structure (Figure 1 a-b). Therefore, the structures determined to date of wild-type and mutant α-syn fibrils demonstrate that the conserved kernel formed by residues 50-77 acts a modular building block to assemble protofilaments into distinct polymorphs in at least four different ways.

In all seven α-syn fibril structures, the conserved kernel features a cavity, possibly a solvent channel surrounded by T54, A56, K58, G73, and V74 (Figure 1 b, Figure 2 c, Supplementary Figure 6 b)^14, 27–29^. However, the H50Q polymorphs determined here are so far the only α-syn fibril structures to resolve density in this cavity. A hydrophilic cavity is also observed in the human serum amyloid A cryo-EM structure and two structures of immunoglobulin light chain fibrils^22, 30, 31^. Therefore, full-length amyloid fibrils share the property of crystal structures of amyloid segments where the majority of the protein packs in a manner excluding water, while some water molecules can be seen hydrogen-bonding with the backbone or polar side chains^32^. Recent structures of tau fibrils extracted from the brains of chronic traumatic encephalopathy patients suggest that a hydrophobic molecule may occupy a hydrophobic cavity present in the fibril core^33^. This is also similar to crystal structures of amyloid segments where molecules such as polyethylene glycol can occupy cavities in the crystal packing^34^.

The structures of H50Q polymorphs help explain the differences in aggregation kinetics, seeding capacity, and cytotoxicity between H50Q and wild-type α-syn we and others have observed (Figure 4 b, d-e)^18, 19, 25^. First, the observation that the H50Q mutation results in a large proportion of Narrow Fibrils formed from a single protofilament may help explain the faster aggregation kinetics of the mutant fibrils. Given that multiple molecules must come together to nucleate amyloid fibril growth, fibrils composed of a single protofilament may therefore have a shorter lag time than fibrils composed of two protofilaments as half as many molecules are required to form the nucleus. Thus, the H50Q mutation may lower the barrier to nucleation and more readily lead to fibril formation. Previous studies on wild-type α-syn have demonstrated that minor species of single protofilament fibrils exist in polymorphic preparations, suggesting that wild-type a-syn can also form single protofilaments^20^. However, here we observe that single protofilament structures dominate, suggesting that the H50Q mutation may tip the balance to favor single protofilament fibrils and shorter lag times.

Second, the higher seeding capacity of H50Q versus wild-type fibrils in our biosensor cell assays may be explained by the presence of a secondary nucleation mechanism. We speculate that the Wide Fibril species represents a step in a secondary nucleation pathway whereby residues ^58^KTKE^61^ of Protofilament A serve as a surface to catalyze the formation of Protofilament B (Supplementary Figure 11)^35^. Several observations support this idea: 1) Protofilament B is never observed alone, and is only observed with Protofilament A in the Wide Fibril whereas Protofilament A is observed alone in the Narrow Fibril. 2) The structure of Protofilament A in both the Narrow and Wide Fibril is nearly identical (R.M.S.D=0.26 Å) and the helical twist of both the Narrow and Wide Fibril is nearly identical (crossover distance ∼900 Å) suggesting that Protofilament A acts as an unperturbed scaffold for Protofilament B to grow off its side. 3) The Wide Fibril protofilament interface is exceedingly small, perhaps making it a labile interface where Protofilament B can nucleate and elongate but eventually fall off, forming individual single protofilament fibrils. Given that we do not observe Protofilament B alone, we speculate that over time Protofilament B may convert into Protofilament A perhaps initiated by the switching of H50Q into a conformation that allows hydrogen bonding with K45 and subsequent formation of the Y39-T44-E46 hydrogen bond triad. We speculate the conversion of Protofilament B to Protofilament A may occur immediately before disassembling from the Wide Fibril to form individual Narrow Fibrils or after disassembly from the Wide Fibril (Supplementary Figure 11).

Third, the ultrastructural arrangement of protofilaments, the ordered β-arch in Protofilament A featuring a unique Y39-T44-E46 hydrogen bond triad, and the presence of Islands 1 and 2 represent major structural differences in the H50Q fibrils compared to previously determined wild-type α-syn structures. These structural differences create new ordered surfaces on the fibrils that may enable their heightened cytotoxicity compared to wild-type. Previously, it has been shown that polyQ inclusions can sequester essential cellular proteins and that re-supply of sequestered proteins ameliorated toxicity and reduced inclusion size, presumably by coating the fibrils and rendering them inert^36^. This highlights the importance of surfaces of fibrils in mediating cytotoxicity and suggests that different surface properties of amyloid fibrils may explain differential cytotoxicites. Indeed, others have shown that two polymorphs of wild-type α-syn could be homogenously prepared and that these polymorphs had different cytotoxicities to SH-SY5Y cells suggesting that differences in structure, including exposed fibril surfaces, result in different cytotoxicity^37^. Therefore, we propose that the large structural differences observed in our H50Q fibrils compared to wild-type could mediate the higher cytotoxicity of H50Q versus wild-type fibrils.

Hereditary mutations in α-syn are largely clustered in the preNAC region (residues 47-56) away from the NACore region (residues 68-78). The E46K hereditary mutation is predicted to disrupt a key salt bridge that forms between E46 and K80, potentially disturbing the fibril core. Consistent with this, NMR studies have shown large chemical shifts for residues in the fibril core of E46K fibrils^38^. Interestingly, in H50Q Protofilament A, E46 participates in a hydrogen bond network with T44 and Y39; therefore, mutation to lysine may disrupt this network making the formation of the N-terminal 36-46 β-arch mutually exclusive with the E46K hereditary mutation. Further, the observation that the E46-K80 hydrogen bond is not maintained in Protofilament A – unlike the previous wild-type rod polymorphs and Protofilament B – demonstrates that this interaction is not necessary to maintain the overall wild-type fold, and that in the case of the E46K hereditary mutation, it is the change to lysine and unfavorable juxtaposition of the positively charged E46K and K80 that explains the rearrangement of the fibril core as indicated by NMR^38^. For mutations A30P and A53T, NMR studies of fibrils show small perturbations in chemical shifts and secondary structures at sites proximal to the mutation, suggesting the overall fold of the fibril is largely unchanged^38, 39^. Here, the H50Q mutation seems to lie somewhere in the middle of the A30P, A53T, and E46K mutations; in that, H50Q enforces a new conformation of the N-terminus of the fibril core, disrupts previously observed protofilament interfaces, and creates a new protofilament interface while maintaining a conserved β-arch fold in the fibril core. Further work is needed to determine the exact structural effects of other α-synuclein hereditary mutations.

Overall, our results demonstrate that the H50Q hereditary mutation leads to new fibril polymorphs that have more rapid fibril-forming kinetics, higher seeding capacity, and higher cytotoxicity. These findings provide a starting point for understanding the structural basis of mutation-enhanced pathogenesis in the synucleinopathies.

## Acknowledgments

We thank H. Zhou for use of Electron Imaging Center for Nanomachines (EICN) resources and P. Ge for assistance in cryo-EM data collection. We acknowledge the use of instruments at the EICN supported by NIH (1S10RR23057 and 1S10OD018111), NSF (DBI-1338135), and CNSI at UCLA. The authors acknowledge NIH AG 054022, NIH AG061847, and DOE DE-FC02-02ER63421 for support. D.R.B. was supported by the National Science Foundation Graduate Research Fellowship Program.

## Author Contributions

D.R.B. and B.L. designed experiments and performed data analysis. B.L. and C.S. expressed and purified the α-syn protein. B.L. grew fibrils of α-syn and performed biochemical experiments. D.R.B. and B.L prepared cryo-EM samples and performed cryo-EM data collection. B.L. and W.F. selected filaments from cryo-EM images. D.R.B. performed cryo-EM data processing and built the atomic models. M.R.S. wrote the software for and D.R.B. carried out solvation energy calculations. All authors analyzed the results and D.R.B wrote the manuscript with input from all authors. L.J. and D.S.E. supervised and guided the project.

## Competing Interests

D.S.E. is an advisor and equity shareholder in ADRx, Inc.

## Supporting Information

### Methods

#### Protein purification

Full-length aSyn WT and H50Q mutant proteins were expressed and purified according to a published protocol^1^. The bacterial induction started at an OD600 of ∼0.6 with 1 mM IPTG for 6 h at 30°C. The harvested bacteria were lysed with a probe sonicator for 10 minutes in an iced water bath. After centrifugation, the soluble fraction was heated in boiling water for 10 minutes and then titrated with HCl to pH 4.5 to remove the unwanted precipitants. After adjusting to neutral pH, the protein was dialyzed overnight against Q Column loading buffer (20 mM Tris-HCl, pH 8.0). On the next day, the protein was loaded onto a HiPrep Q 16/10 column and eluted using elution buffer (20 mM Tris-HCl, 1M NaCl, pH 8.0). The eluent was concentrated using Amicon Ultra-15 centrifugal filters (Millipore Sigma) to ∼5 mL. The concentrated sample was further purified with size-exclusion chromatography through a HiPrep Sephacryl S-75 HR column in 20 mM Tris, pH 8.0. The purified protein was dialyzed against water, concentrated to 3 mg/ml, and stored at 4C. The concentration of the protein was determined using the Pierce™ BCA Protein Assay Kit (cat. No. 23225, Thermo Fisher Scientific).

#### Fibril preparation and optimization

Both WT and H50Q fibrils were grown under the same condition: 300 µM purified monomers, 15mM tetrabutylphosphonium bromide, shaking at 37°C for 2 weeks.

#### Negative stain transmission electron microscopy (TEM)

The fibril sample (3 μL) was spotted onto a freshly glow-discharged carbon-coated electron microscopy grid. After 1 minute, 6 μL uranyl acetate (2% in aqueous solution) was applied to the grid for 1 minutes. The excessive stain was remove by a filter paper. Another 6μL uranyl acetate was applied to the grid and immediately removed. The samples were imaged using an FEI T20 electron microscope.

#### Thioflavin-T binding assay

50 μM of purified aSyn monomers were adequately mixed with 20 μM thioflavin T and added into a 96-well-plate. Samples were incubated at 37°C for 2 days with 600 rpm double orbital shaking. The ThT signal was monitored using the FLUOstar Omega Microplate Reader (BMG Labtech; Cary, NC) at an excitation wavelength of 440 nm and an emission wavelength of 490 nm.

#### SDS Stability

SDS was diluted in water to make SDS solutions at 2.5%, 5%, 10% and 15%. Fibrils at the end of the ThT assay were treated with SDS solution at 5:1 volume ratio to obtain SDS concentrations of 0.5%, 1%, 2% and 3%. Each solution was transferred to 3 microcentrifuge tubes and heated at 70° C for 15 minutes. After the treatment, the ThT signal was obtained. The 0% SDS solution without heating was treated with equal amount of water and used for normalization.

#### Fibril seeding aggregation in cells

We performed the biosensor cell seeding assay based on a previously published protocol^2^. Briefly, the assay works as follows: exogenous, un-labeled fibrils are transfected into HEK293T cells expressing α-syn-A53T-YFP. Seeded aggregation of endogenously expressed α-syn-A53T-YFP is monitored by formation of fluorescent puncta. The puncta represent a condensation of α-syn-A53T-YFP as a result of seeding by exogenous H50Q or WT fibrils.

Human embryonic kidney FRET Biosensor HEK293T cells expressing full-length aSyn containing the hereditary A53T mutation were grown in DMEM (4mM L-glutamine and 25mM D-glucose) supplemented with 10% FBS, 1% penicillin/streptomycin. Trypsin-treated HEK293T cells were harvested, seeded on flat 96-well plates at a concentration of 4×104 cells/well in 200 μL culture medium per well and incubated in 5% CO2 at 37°C for 18 hours.

aSyn fibrils were prepared by diluting with Opti-MEM™ (Life Technologies; Carlsbad CA) and sonicating in a water bath sonicator for 10 minutes. The fibril samples were then mixed with Lipofectamine™ 2000 (Thermo Fisher Scientific) and incubated for 15 minutes and then added to the cells. The actual volume of Lipofectamine™ 2000 was calculated based on the dose of 1 μL per well. After 48 hours of transfection, the cells were trypsinized, transferred to a 96-well round-bottom plate and resuspended in 200 μL chilled flow cytometry buffer (HBSS, 1% FBS, and 1 mM EDTA) containing 2% paraformaldehyde. The plate was sealed with Parafilm and stored at 4 °C for imaging. Fluorescent images were processed in ImageJ to count number of seeded cells (Supplementary Figure 12).

#### MTT mitochondrial activity assay

The addition of sonicated fibrils to Nerve Growth Factor-differentiated PC12 cells is a well-established assay to measure cytotoxicity of amyloid fibrils^3, 4^. Use of this neuron-like cell line allows us to obtain a biologically relevant assay for cytotoxicity. For our MTT mitochondrial activity assay, the protocol was adapted from the Provost and Wallert laboratories^5^. Thiazolyl blue tetrazolium bromide for the MTT cell toxicity assay was purchased from Millipore Sigma (M2128-1G; Burlington, MA). PC12 cells were plated in 96-well plates at 10,000 cells/well in DMEM (Dulbecco’s modification of Eagle’s medium; 5% fetal bovine serum [FBS], 5% heat-inactivated horse serum, 1% penicillin/streptomycin and 150 ng/mL nerve growth factor 2.5S (Thermo Fisher Scientific). The cells were incubated for 2 days in an incubator with 5% CO2 at 37°C. The cells were treated with different concentrations of aSyn fibrils (200 nM, 500 nM,1000 nM, 2000 nM). After 18 hours of incubation, 20 μl of 5 mg/ml MTT was added to every well and the plate was returned to the incubator for 3.5 hours. With the presence of MTT, the experiment was conducted in a laminar flow hood with the lights off and the plate was wrapped in aluminum foil. The media was then removed with an aspirator and the remained formazan crystals in each well were dissolved with 100 μL of 100% DMSO. Absorbance was measured at 570 nm to determine the MTT signal and at 630 nm to determine background. The data were normalized to those from cells treated with 1% SDS to obtain a value of 0%, and to those from cells treated with PBS to obtain a value of 100%.

#### Lactate Dehydrogenase (LDH) Assay

The LDH viability assay was performed using the CytoTox-ONE™ Homogeneous Membrane Integrity Assay (Promega; Madison, WI, G7891). PC12 cells were cultured and differentiated with the same protocol as described in the MTT assay. Different concentrations of aSyn fibrils (200 nM, 500 nM, 1000 nM, 2000nM) were added to the cells for 18 hours of incubation with 5% CO2 at 37°C. The assay was carried out in a 96 well plate and the fluorescent readings were taken in the FLUOstar Omega Microplate Reader (Ex. 560 nm, Em. 590 nm, BMGLabtech; Cary, NC). PBS and 0.2% Triton-X100 treated cells were used as negative and positive control for normalization.

#### Cryo-EM data collection and processing

2 µl of fibril solution was applied to a baked and glow-discharged Quantifoil 1.2/1.3 electron microscope grid and plunge-frozen into liquid ethane using a Vitrobot Mark IV (FEI). Data were collected on a Titan Krios (FEI) microscope equipped with a Gatan Quantum LS/K2 Summit direct electron detection camera (operated with 300 kV acceleration voltage and slit width of 20 eV). Counting mode movies were collected on a Gatan K2 Summit direct electron detector with a nominal physical pixel size of 1.07 Å/pixel with a dose per frame 1.2 e-/Å^2^. A total of 30 frames with a frame rate of 5 Hz were taken for each movie resulting in a final dose 36 e-/Å^2^ per image. Automated data collection was driven by the Leginon automation software package^6^.

Micrographs containing crystalline ice were used to estimate the anisotropic magnification distortion using mag_distortion_estimate^7^. CTF estimation was performed using CTFFIND 4.1.8 on movie stacks with a grouping of 3 frames and correction for anisotropic magnification distortion^8^. Unblur^9^ was used to correct beam-induced motion with dose weighting and anisotropic magnification correction, resulting in a physical pixel size of 1.065 Å/pixel.

All particle picking was performed manually using EMAN2 e2helixboxer.py^10^. We manually picked two groups of particles for further data processing: the first group was composed of all fibrils and the second group was composed of Wide Fibrils. For the first group, particles were extracted in RELION using the 90% overlap scheme into 1024 and 288 pixel boxes. Classification, helical reconstruction, and 3D refinement were used in RELION as described^11^. For the first group of all particles, we isolated Narrow Fibrils during 2D Classification and subsequently processed them as a separate data set. 2D Classifications of Narrow Fibril 1024 pixel boxes were used to estimate helical parameters. We performed 3D classification with the estimated helical parameters for Narrow Fibrils and an elongated Gaussian blob as an initial model to generate starting reconstructions. We ran additional 3D classifications using the preliminary reconstructions from the previous step to select for particles contributing to homogenous classes (stable helicity and separation of β-strands in the X-Y plane). Typically, we performed Class3D jobs with K=3 and manual control of the tau_fudge factor and healpix to reach a resolution of ∼5-6 Å to select for particles that contributed to the highest resolution class for each structure. We employed Refine3D on a final subset of Narrow Fibril particles with 288 pixel box size to obtain the final reconstruction. We performed the map-map FSC with a generous, soft-edged solvent mask and high-resolution noise substitution in RELION PostProcess resulting in a resolution estimate of 3.3 Å.

We extracted particles from the Wide Fibril data set using 1024 and 686 pixel boxes. 2D Classifications of 1024 and 686 pixel boxes were used to estimate helical parameters. 2D Classifications of 686 pixel boxes were used to further isolate only Wide Fibril segments since there was still some other fibril species that were included due to the fact that we could not separate all fibril species perfectly during manual picking. Once a homogenous set of Wide Fibrils was obtained during 2D Classification of 686 pixel boxes, we performed a 3D reconstruction using an elongated Gaussian blob as an initial model. The asymmetry present in the 686 box 2D class averages of the Wide Fibril (Supplementary Figure 3 c) prompted us to use a helical rise of 4.8 Å and C1 symmetry due to the fact that if a 2-fold symmetry were present in the fibrils, 2D class averages would display a mirror symmetry across the fibril axis. After an initial 2D model was generated for the Wide Fibril, we re-extracted all tubes corresponding to those particles included in the final subset of Wide Fibril 686 pixel boxes with a box size of 224 pixels. All 224 pixel boxes were subjected to multiple rounds of 3D Classification using the initial 686 pixel box Wide Fibril reconstruction as a reference. We refined the final subset of particles using Refine3D to a resolution of 3.6 Å. We performed resolution estimation as described above for the Narrow Fibrils.

#### Atomic model building

We sharpened both the Narrow and Wide Fibril reconstructions using phenix.auto_sharpen^12^ at the resolution cutoff indicated by the map-map FSC and subsequently built atomic models in to the refined maps with COOT^13^. We built the model for the Narrow Fibril *de novo* using previous structures of wild-type alphα-synuclein fibrils as guides. To build the Wide Fibril model, we made a copy of one chain of the Narrow Fibril structure and rigid-body fit it into the second protofilament density observed in the Wide Fibril reconstruction. This resulted in the Wide Fibril being composed of one protofilament nearly identical to the Narrow Fibril and one protofilament with less ordered N- and C-termini, therefore resulting in an asymmetric double protofilament structure.

For both Narrow and Wide Fibrils, we generated a 5-layer model to maintain local contacts between chains in the fibril during structure refinement. We performed automated structure refinement for both Narrow and Wide Fibrils using phenix.real_space_refine^14^. We employed hydrogen bond distance and angle restraints for backbone atoms participating in β-sheets and side chain hydrogen bonds during automated refinements. We performed comprehensive structure validation of all our final models in Phenix.

Although we did not include coordinates in our final models for additional residues that could occupy Islands 1 and 2 neighboring Protofilament A because we could not be certain which residues occupy those densities, we built several speculative models (Supplementary Figure 4). For Island 2, we assumed that there was a short disordered linker between residue 36 of Protofilament A and Island 2 resulting in the residues occupying Island 2 forming a tight interface with ^36^GVLYVG^41^ of the fibril core. We noticed that the sequence ^32^KTKE^35^ immediately precedes the last ordered residue of Protofilament A, G36. ^32^KTKE^35^ often forms the bends that connect straight β-strands in the ordered fibril core, so we assumed that this sequence would be a good candidate to form the tight bend that connects G36 to Island 2. Therefore, we modeled in residues ^26^VAEAAG^31^ into Island 2. These residues satisfied the requirements of having short, hydrophobic side chains forming the tight interface with residues V38 and L40 from the fibril core.

For Island 1, we assumed the densities either come from the N-terminus of Protofilament A or the C-terminus of Protofilament B. For the former case, we assume that there is a minimum of ∼8 residues that are disordered between the end of Island 2 and the beginning of Island 1 (see Supplementary Figure 4). This is because there is 27 Å between the end of Island 2 - V26 - and the beginning of Island 1, and we assume a minimum of ∼3.3 Å/residue. Therefore, we threaded 8 residues at a time from the region ^1^MDVFMKGLSKAKEGVVAAA^20^ onto a β-strand backbone placed in the Island 1 density to identify candidate 8mers. While most sequences could not plausibly occupy Island 1 due to steric clashes with the preNAC region of the fibril core, or due to glycine residues occupying positions where there were obvious side chain densities, several candidate sequences were identified (Supplementary Figure 4).

We followed a similar protocol to identify possible sequences from the C-terminus. In this case, we assume that these residues could come from either a largely disordered Protofilament B molecule in the Narrow Fibril or an ordered Protofilament B molecule as seen in the Wide Fibril (see Supplementary Figure 4). Here, we assume there is a minimum of ∼15 residues from the last ordered residue of the C-terminus of Protofilament B - since we assume a minimum of ∼3.3 Å/residue and there is ∼45 Å between K97 of Protofilament B and the beginning of Island 1 (see Supplementary Figure 4). Therefore, we threaded all possible 8mers from the region ^112^ILEDMPVDPDNEAYEMPSEEGYQDYEPEA^140^ onto a β-strand backbone occupying Island 1 to identify candidate sequences following the same criterion as above.

We created the speculative model of an H50Q double protofilament containing a homomeric preNAC interface by aligning a single chain from the H50Q Narrow Protofilament with the helical axis of the wild-type “rod” structure (6CU7) and applying a pseudo-2(1) helical symmetry to generate a symmetrically related second chain.

#### Energetic calculation

The stabilization energy is an adaptation of the solvation free energy described previously^15^, in which the energy is calculated as the sum of products of the area buried of each atom and its corresponding atomic solvation parameter (ASP). ASPs were taken from our previous work^15^. Area buried is calculated as the difference in solvent accessible surface area (SASA) of the reference state (i.e. unfolded state) and the SASA of the folded state. The reference state was measured absent all other atoms in the structure but the residue “i” and main chain atoms of residue i-1 and i+1. The SASA of the folded state was measured for each atom in the context of all amyloid fibril atoms. Fibril coordinates were extended by symmetry by three to five chains on either side of the reported molecule, to ensure the energetic calculations were representative of the majority of molecules in a fibril, rather than a fibril end. To account for energetic stabilization of main chain hydrogen bonds, the ASP for backbone N/O elements was reassigned from −9 to 0 if they participated in a hydrogen bond. Similarly, if an asparagine or glutamine side chain participated in a polar ladder (two hydrogen bonds per amide), and was shielded from solvent (SASAfolded < 5 Å^2^), the ASPs of the side chain N and O elements were reassigned from −9 to 0. Lastly, the ASP of ionizable atoms (e.g. Asp, Glu, Lys, His, Arg, N-terminal amine, or C-terminal carboxylate) were assigned the charged value (−37/-38) unless the atoms participated in a buried ion pair, defined as a pair of complementary ionizable atoms within 4.2 Å distance of each other, each with SASAfolded < 40 Å^2^). In that case, the ASP of the ion pair was reassigned to −9. In the energy diagrams, a single color is assigned to each residue, rather than each atom. The color corresponds to the sum of solvation free energy values of each of the atoms in the residue. The energy reported for FUS in Table 2 is the average over 20 NMR models. The standard deviation is 1.8 kcal/mol.

### Supplementary Note(s)

#### Supplementary Note 1: Un-modeled Density Flanking Protofilament A

In Protofilament A in both the Narrow and Wide Fibril polymorphs, there are two regions of unmodeled density resembling β-strands that flank residues Q50-E57 and G36-V40 that we term Island 1 and Island 2, respectively (Figure 1 c, Supplementary Figure 4). Since we cannot unambiguously assign residues to these densities, we have made several speculative molecular models for the residues in the Islands (Supplementary Figure 4). First, for Island 2, we assume that there is likely a short, disordered region that precedes the first ordered residue of the fibril core, G36, and follows the first residue in Island 2. We noticed that G36 is preceded by sequence 32KTKE35, a repeated sequence motif in α-synuclein that forms sharp turns in other segments of the fibril. Therefore, we assume that 32KTKE35 may form a similar turn that links Island 2 to G36, and we thus modeled residues 26VAEAAG31 into Island 2 (Supplementary Figure 4 c). These residues satisfy the constraint of maintaining a tight, hydrophobic interface with residues L38 and V40 of the fibril core and demonstrate a reasonable fit into the Island 2 electron density (Supplementary Figure 4 c).

Assigning residues to Island 1 is more ambiguous as there are no close-by termini that could result in a short disordered linker region before the chain becomes ordered again at Island 1, as we believe is the case for Island 2. Therefore, we reasoned that either the residues occupying Island 1 come from Protofilament A or B (Supplementary Figure 4 a, b) or from a different chain of α-syn. The latter case would support the idea of secondary nucleation, whereby the side of an amyloid fibril provides a template for additional molecules to become ordered and eventually form mature amyloid fibrils^16^. However, since there are too many possible residues within the entire α-syn sequence that could occupy Island 1, we decided to first make models for sequences that could come from the N-terminus of Protofilament A (in either the Narrow or Wide Fibril) or the C-terminus of Protofilament B. This is due to the fact that these two termini are positioned close enough to Island 1 to allow residues from the N- or C-terminus to account for the Island 1 density (Supplementary Figure 4 a-b, see Methods).

We identified several possible sequences from the N- or C-terminus that could account for Island 1 in Protofilament A (Supplementary Figure 4 c, see Methods). We also show several examples of sequences that were eliminated from consideration due to steric clashes, glycine residues at positions where there was obvious side chain density, or proline residues in the middle of a β-strand (Supplementary Figure 4 c). Candidate sequences include 3VFMKGLSK10, 7GLSKAKEG14, and 10KAKEGVVA17 from the N-terminus and 117PDNEAYEM124 from the C-terminus (Supplementary Figure 4 c).

We also noticed that Island 1 is located in the same region as the protofilament interface of the rod polymorph^1^, perhaps suggesting that a portion of another protein chain is interacting with the preNAC residues from Protofilament A. We therefore modeled in residues from the preNAC into Island 1 in a manner similar to the wild-type rod polymorph (Supplementary Figure 5 a)8. We find that residues 50QGVATVA56 match the Island 1 density well and form a tight steric zipper with the preNAC residues on Protofilament A (Supplementary Figure 5 a). Energetic analysis of a model with Island 1 as residues 50QGVATVA56 and Island 2 as residues 26VAEAAG31 reveals that Protofilament A is stabilized by the presence of residues in these unmodeled densities (71.6 kcal/mol/layer with islands occupied vs. 58 kcal/mol/layer with islands empty) (Supplementary Figure 5 b). In summary, although we cannot assign unambiguously the residues that occupy Islands 1 and 2, speculative models demonstrate sequences within α-syn that can account for these densities and that the presence of these sequences in Islands 1 and 2 stabilizes Protofilament A.

#### Supplementary Note 2: Speculative models of inter-protofilament interfaces at the H50Q preNAC region

To confirm that H50Q disallows assembly of protofilaments at the preNAC, we made a speculative model of two chains from Protofilament A forming a steric zipper at the preNAC, similar to the wild-type rod polymorphs (Supplementary Figure 8 c). We observe that although the wild-type residues G51-V55 form a tight homomeric steric zipper at the preNAC, H50Q clashes with E57, disfavoring the use of the preNAC as an inter-protofilament interface (Supplementary Figure 8 c). We performed the same analysis in Protofilament B where Q50 is not hydrogen bonding with K45 to examine why we do not observe a protofilament interface formed at the preNAC there. A hypothetical model where the preNAC residues of Protofilament B are used as a protofilament interface reveals that the conformation of Q50 even more severely clashes with E57 on the opposite protofilament than in the hypothetical Protofilament A model and that major clashes occur in other areas as well (Supplementary Figure 8 d). Together, these models help to explain why the preNAC region is disfavored as a protofilament interface in both Protofilaments A and B. Additionally, the severity of the clashes in the Protofilament B model may explain why Protofilament B does not show un-modeled density flanking the preNAC region like Island 2 in Protofilament A (Figure 2 c, Supplementary Figure 8 d).

#### Supplementary Note 3: Differences in stabilization energies for some residues in Protofilament A and B in the Wide Fibril

The differences in stabilizing energies of some residues in the two protofilaments of the Wide Fibril may be due to differences in side chain rotamers that subsequently affect each side chain’s hydrogen bonding and area buried. This is due to the asymmetry in the reconstruction whereby the two protofilaments have slightly different density maps and hence slightly different atomic models, even in the conserved kernel region. For instance, K58 and E61 have different energies in the two protofilaments of the Wide Fibril because they display slightly different hydrogen bonding patterns (Figure 4 a). Since the solvation energy of each residue is the sum of the product of the buried surface area multiplied by the atomic solvation parameter (ASP) for each of its constituent atoms and the ASP for charged atoms depends on if they are involved in a hydrogen bond, the energies for Lys58 and Glu61 can vary depending on their hydrogen bond arrangements. In other cases, such as Glu83 which is not involved in a hydrogen bond network, variation occurs between the two protofilaments because Glu83 adopts two different rotamers. The rotamer with more buried surface area will have a greater stabilization energy.

## Supplementary Figures

**Supplementary Figure 1.**
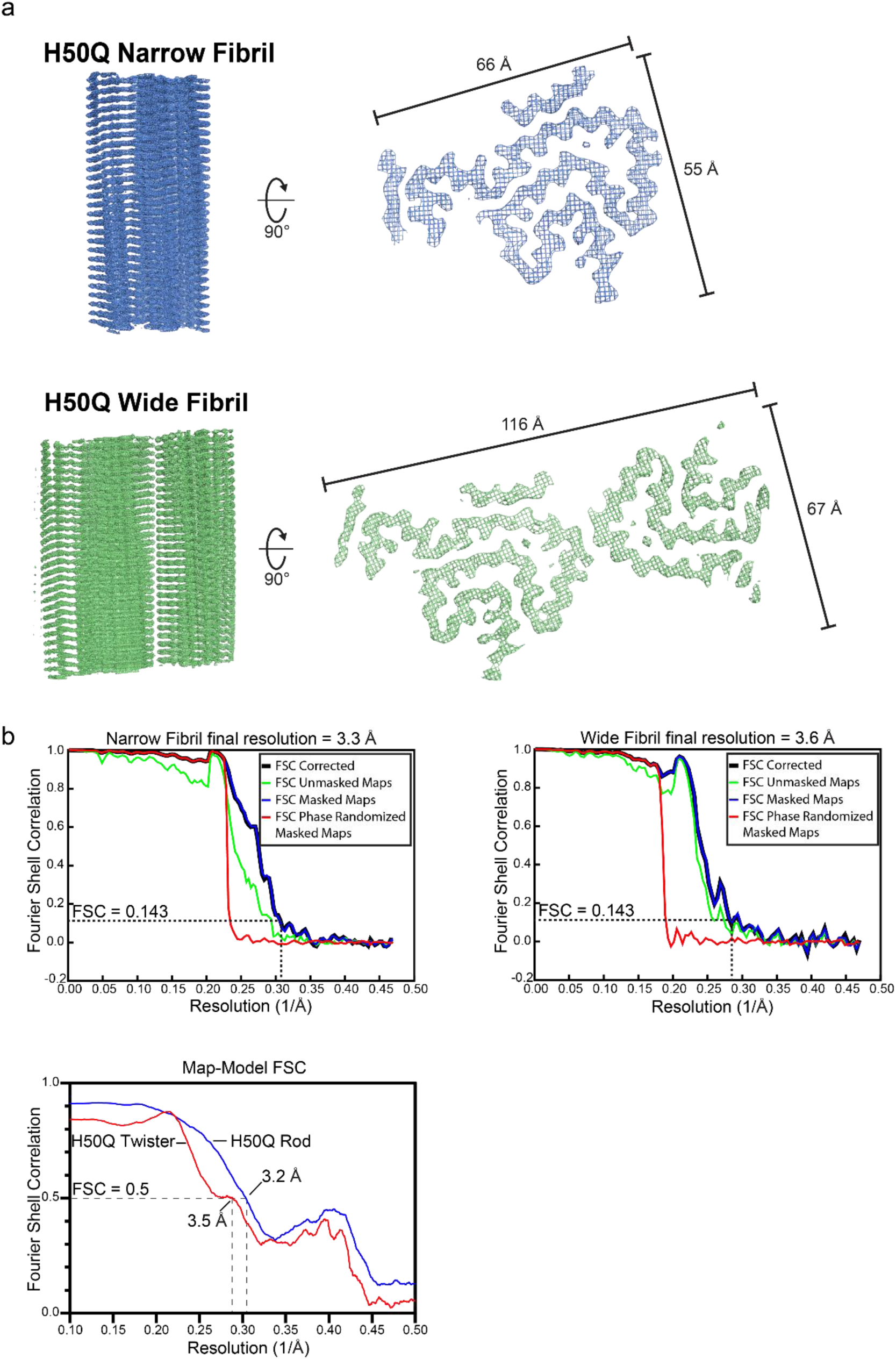
Fourier Shell Analysis a) Helical reconstructions of Narrow and Wide Fibrils with minimum and maximum widths labeled. b) Gold-standard half map FSC curves for Narrow (top, left) and Wide (top, right) Fibrils. Map-model FSC curve for Narrow and Wide Fibrils (bottom).

**Supplementary Figure 2.**
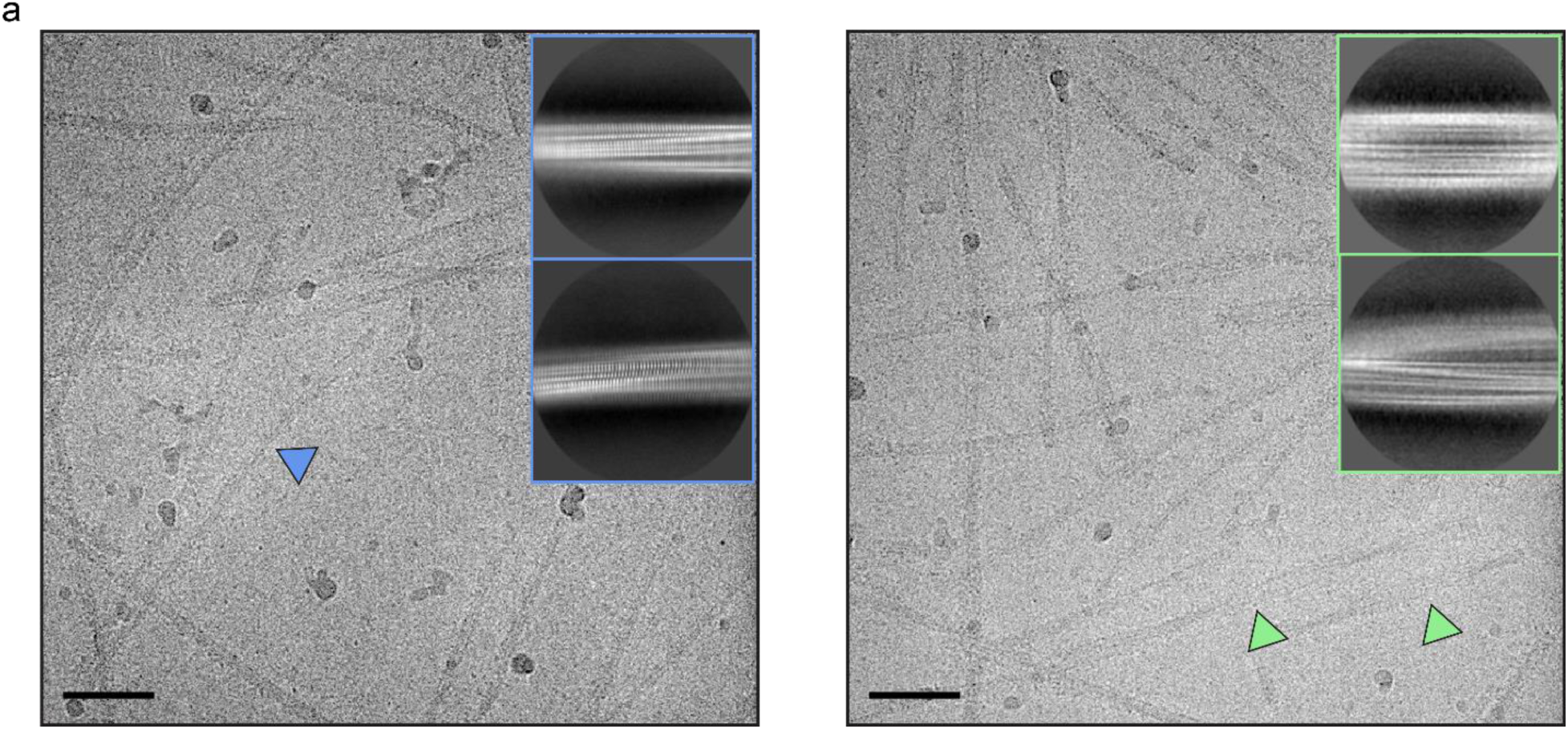
Cryo-EM micrographs and 2D class averages of Narrow (left) and Wide (right) Fibrils. Scale bar = 50 nm.

**Supplementary Figure 3.**
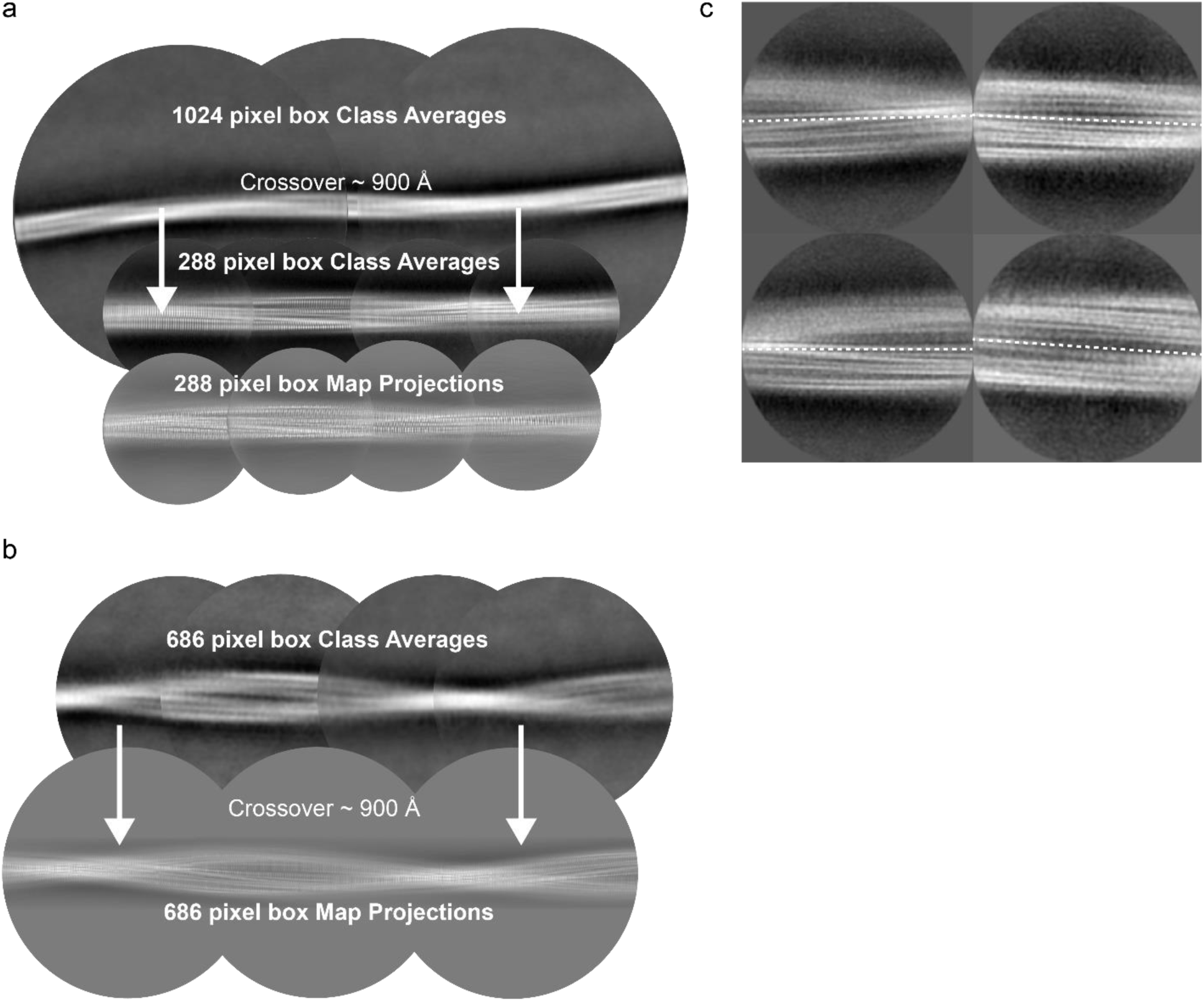
2D class averages and 2D map projections of Narrow and Wide Fibrils. a) 1024 and 288 pixel box size class averages of the Narrow Fibril used to determine crossover distance. 288 pixel box map projections match 2D class averages. b) 686 pixel box size class averages used to determine crossover distance. 686 pixel box map projections match 2D class averages. c) Wide Fibril class averages with a 320 pixel box demonstrate a lack of two-fold symmetry across the fibril axis.

**Supplementary Figure 4.**
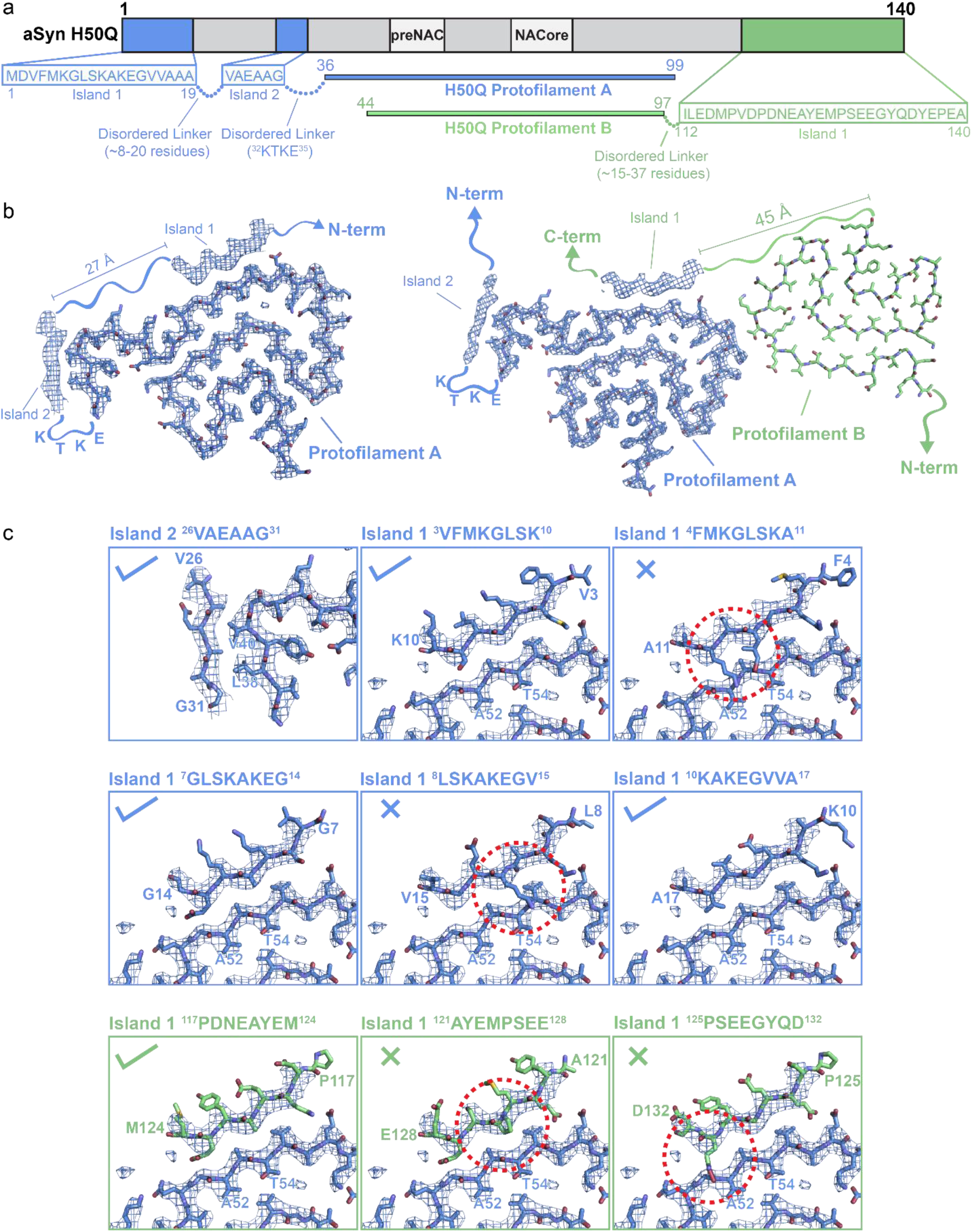
Speculative Atomic Models for Islands 1 and 2 a) Schematic illustrating possible sequences occupying Islands 1 and 2. 8mers from residues 1-19 and 112-140 were considered as possibilities to occupy Island 1. Island 2 is considered to consist of residues 26-31 followed by a disordered linker formed by residues ^32^KTKE^35^. b) Illustration of possible regions from either Protofilament A (left) or Protofilament B (right) that could occupy Island 1. Note that residues from the N-terminus of Protofilament A could account for Island 1 in both the Narrow and Wide Fibril; however, only the Narrow Fibril model is shown here. Island 2 is thought to be formed by the N-terminus of Protofilament A in both Narrow and Wide Fibrils. c) Speculative models for Islands 1 and 2. Check marks indicate plausible models while X’s indicate implausible models. Island 1 models are from either the N-terminus of Protofilament A (blue panels) or the C-terminus of Protofilament B (green panels). Examples of sequences that were found to not be allowed to occupy Island 1 are shown with red dashed circles highlighting steric clashes or β-strand breaking proline residues.

**Supplementary Figure 5.**
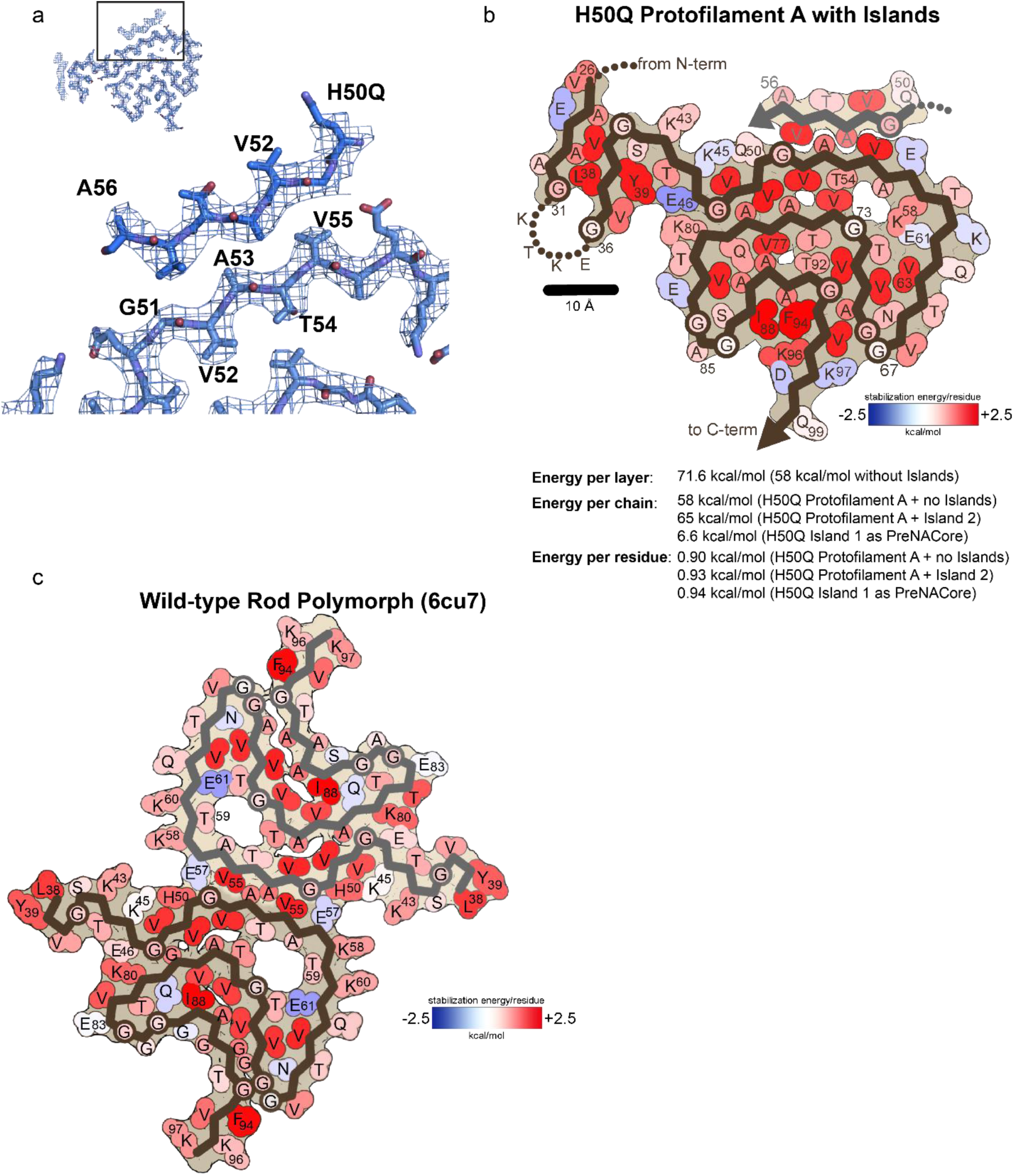
PreNAC homozipper Island 1 model and additional solvation energy maps. a) Speculative model of preNAC residues ^50^QGVATVA^56^ occupying Island 1 in Protofilament A. b) Atomic solvation map and energetic calculations for Protofilament A with Island 1 as ^50^QGVATVA^56^ and Island 2 as ^26^VAEAAG^31^. c) Atomic energy solvation map for Wild-type rod polymorph (6cu7). Notice that K58 and T59 can have favorable stabilization energies whether they are facing the solvent or facing the cavity in the β-arch.

**Supplementary Figure 6.**
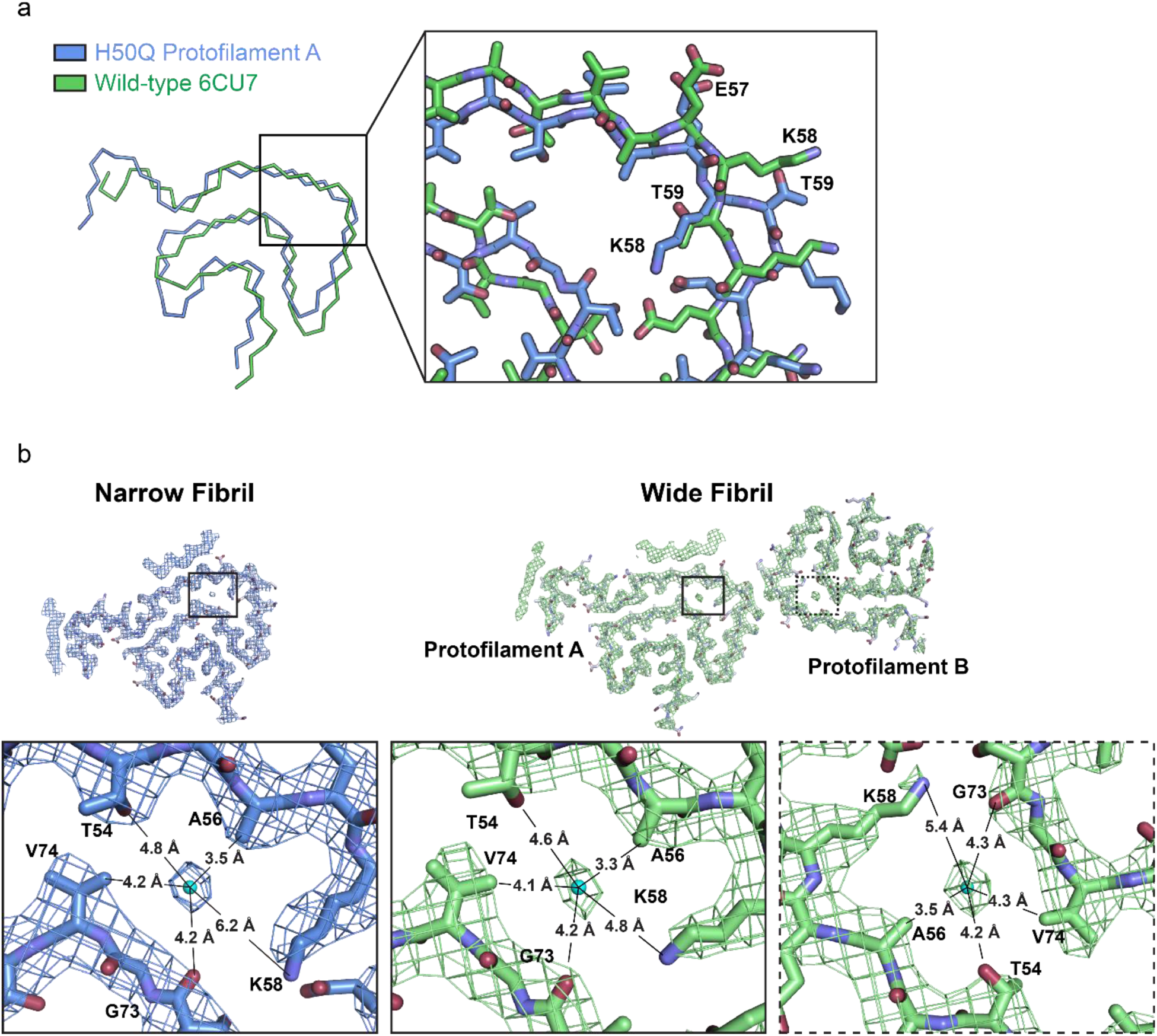
Alternate conformations of K58 and T59 and potential solvent molecules in the α-syn β-arch cavity. a) Wild-type and H50Q fibrils display alternate conformations of K58 and T59. We note that in order for the Wide Fibril to form, T59 needs to be facing away from the fibril core. Therefore the formation of the Wide Fibril is mutually exclusive with our wild-type rod polymorph. b) Environmental distances of putative water molecule for Protofilament A and B in Wide Fibril and b) Protofilament A in Narrow Fibril.

**Supplementary Figure 7.**
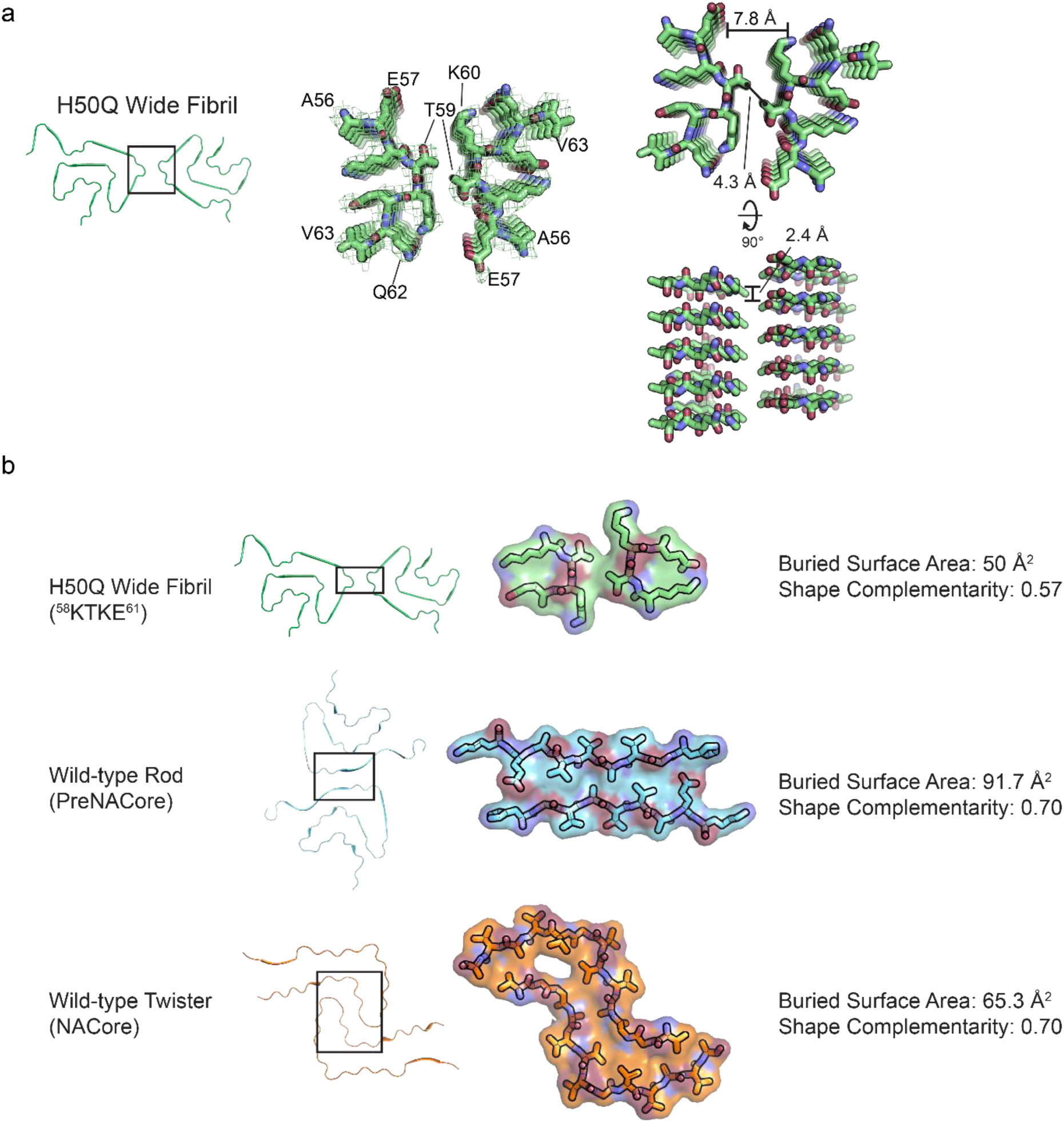
Comparison of α-syn protofilament interfaces a) Wide Fibril overview (left). ^56^AEKTKEQV^63^ homointerface with Wide Fibril electron density (middle). ^56^AEKTKEQV^63^ homointerface showing a 2.4 Å rise between mated strands from Protofilament A and Protofilament B and a distance of 7.8 Å between mated sheets of Protofilament A and B (right). b) Van der Waal’s surface, buried surface area, and shape complementarity of ^58^KTKE^61^ homointerface, preNAC interface, and NACore interface.

**Supplementary Figure 8.**
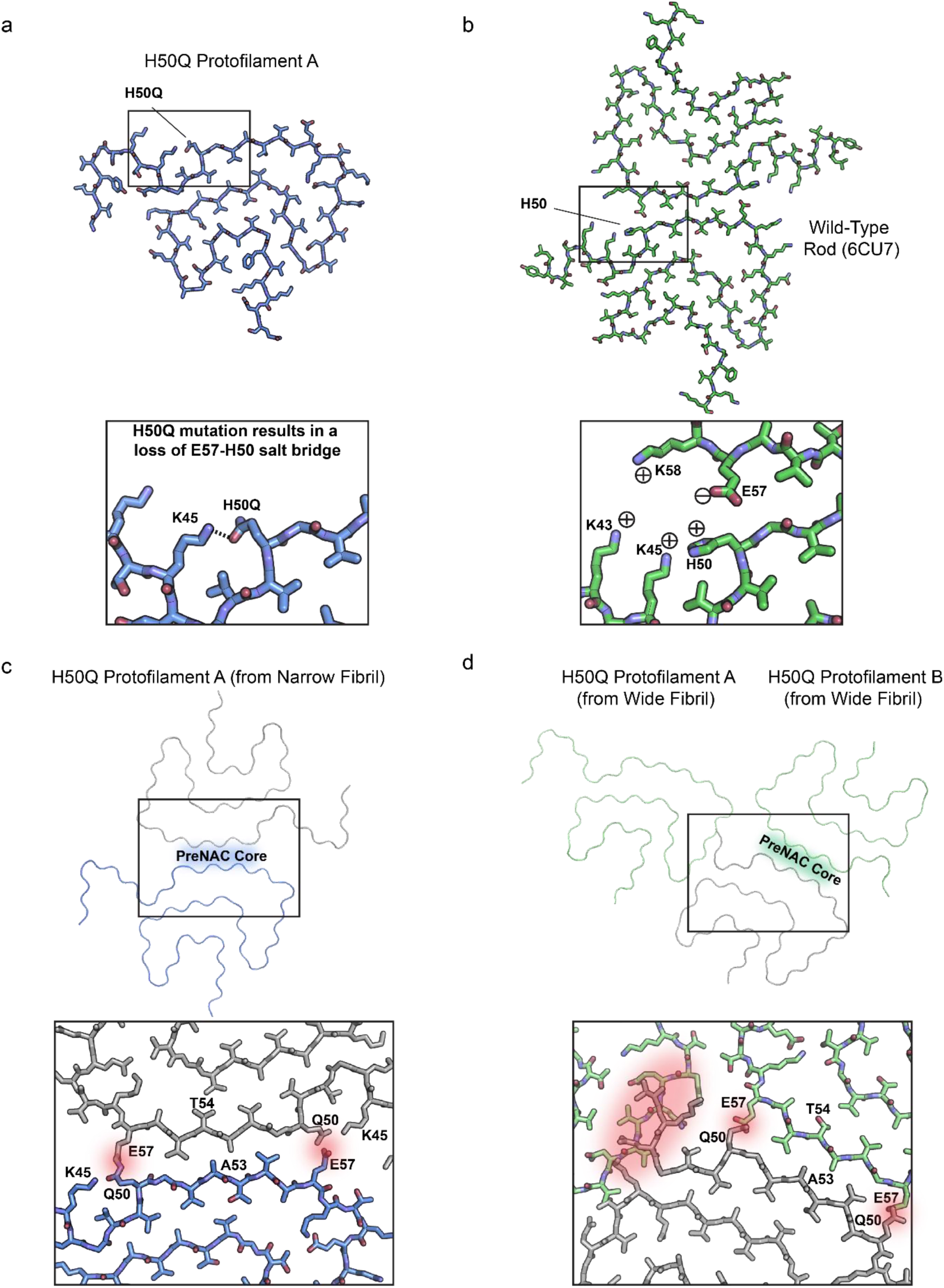
H50Q disrupts the wild-type rod polymorph preNAC protofilament interface a) Conformation of H50Q Protofilament A K45 and H50Q. b) Interaction of K45-H50-E57 in the wild-type rod polymorph protofilament interface. c) Hypothetical H50Q double protofilament using the preNAC of Protofilament A as a steric zipper interface. Notice that the H50Q mutation disfavors the protofilament interface due to steric clashes with E57. d) Hypothetical H50Q protofilament interface using preNAC of Protofilament B. Notice the steric clashes between H50Q and E57 at the hypothetical protofilament interface as well as clashes of other part of the protofilament with Protofilament A.

**Supplementary Figure 9.**
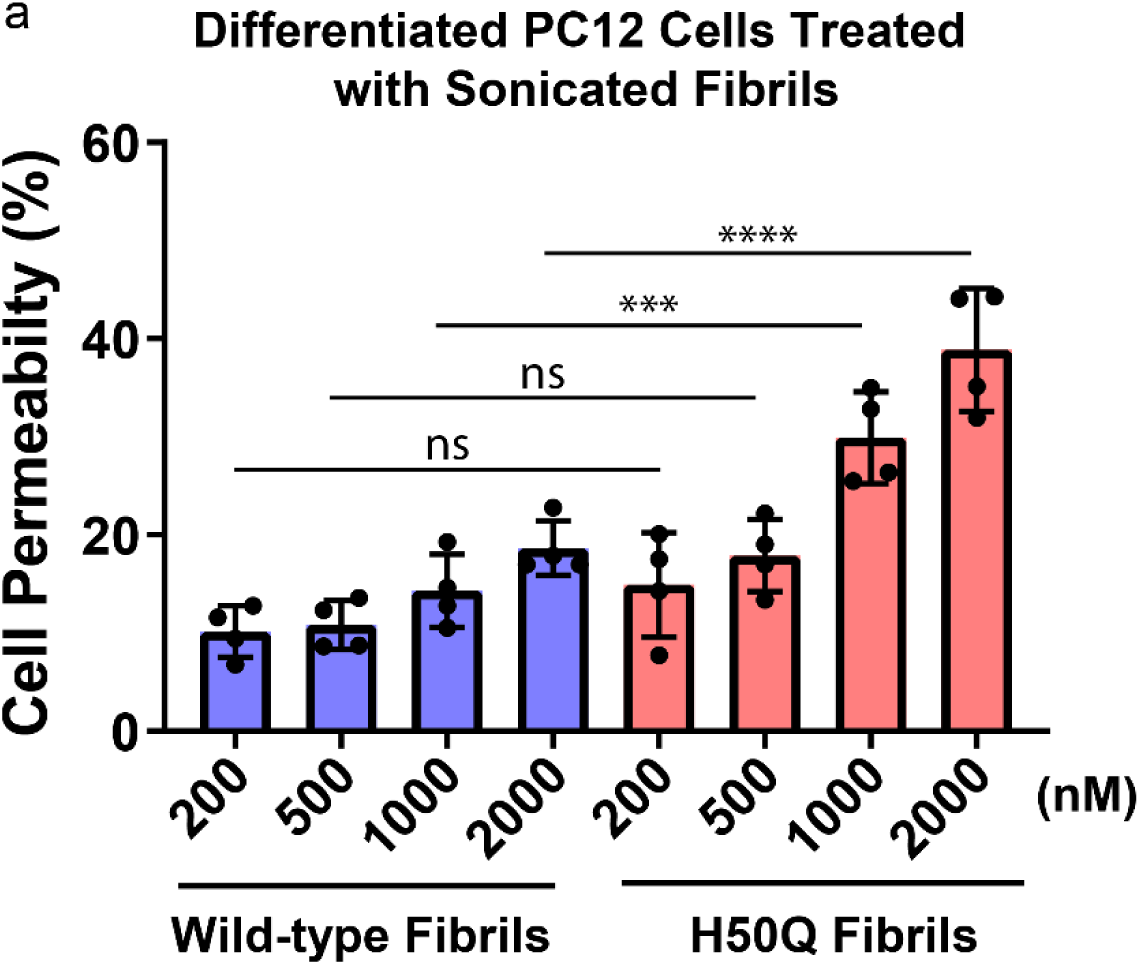
H50Q fibrils disrupt PC12 cell membranes more than WT fibrils. a) Differentiated PC12 cells were treated with sonicated WT and H50Q fibrils and cell permeability was measured via LDH activity in the media (see Methods). H50Q leads to significantly higher cell permeabilization at 1000 and 2000 nM than WT a-syn. Error bars represent standard deviation of four independent experiments. **** = p-value ≤ 0.0001. *** = p-value ≤ 0.001. ns = p-value > 0.05. P-values were calculated using an unpaired, two-tailed t-test with a 95% CI.

**Supplementary Figure 10.**
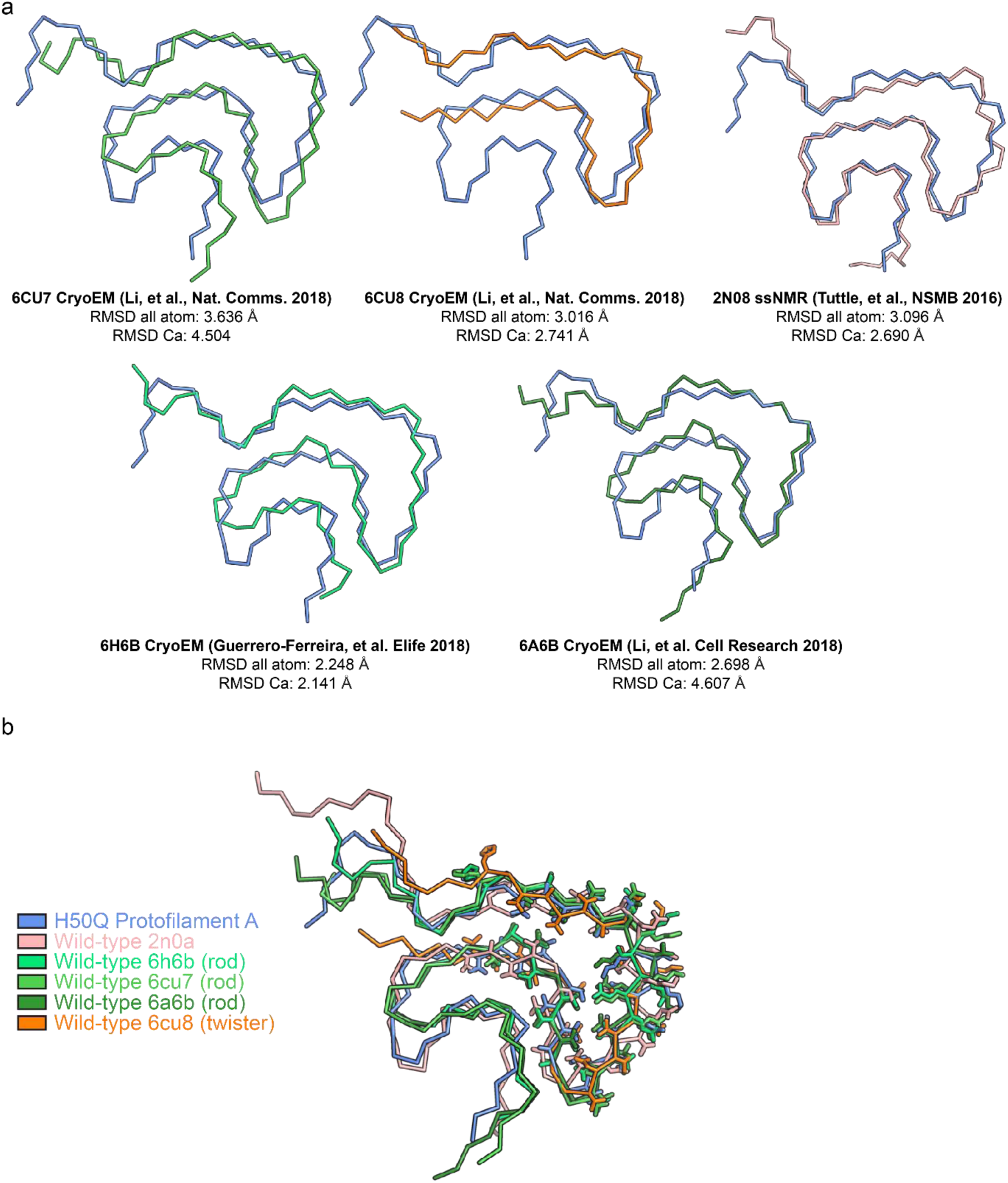
Structural alignment of different wild-type and mutant α-syn polymorphs.

a) Structural alignment of H50Q Protofilament A with all wild-type structures determined thus far. b) Structural alignment of residues 50-57 in wild-type and mutant α-syn polymorphs reveals the kernel region is largely conserved while tail regions, especially the N-terminus adopt variable conformations.

**Supplementary Figure 11.**
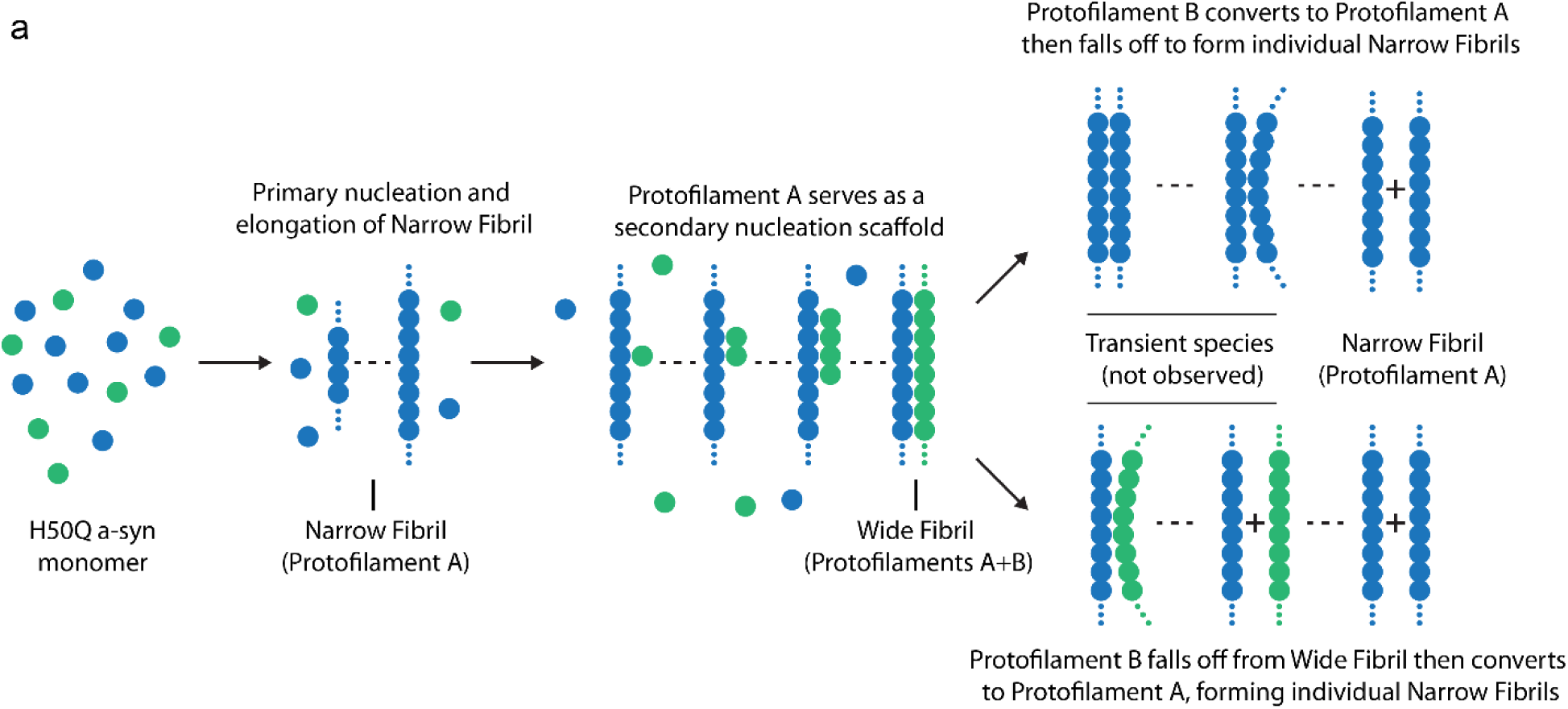
Schematic illustrating possible secondary nucleation of Protofilament B by Narrow Fibrils.

**Supplementary Figure 12.**
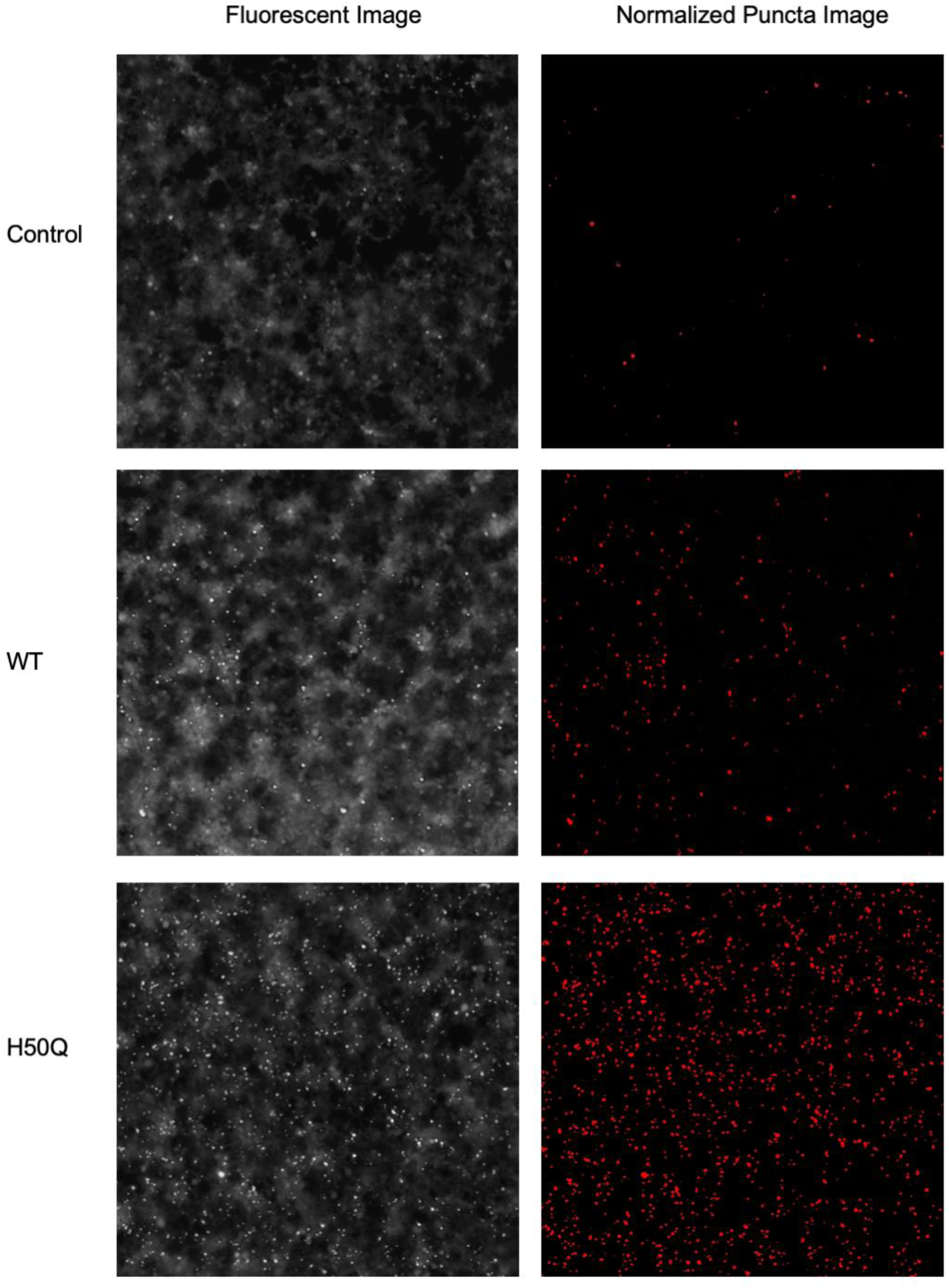
Fluorescent images (left) are taken after seeding aSyn Biosensor cells with 200 nM of sonicated WT and H50Q fibrils (ex. 480 nm, em 580 nm). The images were processed in ImageJ to filter out signal below threshold based on the control’s diffused fluorescent background. Only the individual strong fluorescent puncta are counted as cellular aggregation event (red dots, right).

